# Comparative host interactomes of the SARS-CoV-2 nonstructural protein 3 and human coronavirus homologs

**DOI:** 10.1101/2021.03.08.434440

**Authors:** Katherine M. Almasy, Jonathan P. Davies, Lars Plate

## Abstract

Human coronaviruses have become an increasing threat to global health; three highly pathogenic strains have emerged since the early 2000s, including most recently SARS-CoV-2, the cause of COVID-19. A better understanding of the molecular mechanisms of coronavirus pathogenesis is needed, including how these highly virulent strains differ from those that cause milder, common-cold like disease. While significant progress has been made in understanding how SARS-CoV-2 proteins interact with the host cell, non-structural protein 3 (nsp3) has largely been omitted from the analyses. Nsp3 is a viral protease with important roles in viral protein biogenesis, replication complex formation, and modulation of host ubiquitinylation and ISGylation. Herein, we use affinity purification-mass spectrometry to study the host-viral protein-protein interactome of nsp3 from five coronavirus strains: pathogenic strains SARS-CoV-2, SARS-CoV, and MERS-CoV; and endemic common-cold strains hCoV-229E and hCoV-OC43. We divide each nsp3 into three fragments and use tandem mass tag technology to directly compare the interactors across the five strains for each fragment. We find that few interactors are common across all variants for a particular fragment, but we identify shared patterns between select variants, such as ribosomal proteins enriched in the N-terminal fragment (nsp3.1) dataset for SARS-CoV-2 and SARS-CoV. We also identify unique biological processes enriched for individual homologs, for instance nuclear protein important for the middle fragment of hCoV-229E, as well as ribosome biogenesis of the MERS nsp3.2 homolog. Lastly, we further investigate the interaction of the SARS-CoV-2 nsp3 N-terminal fragment with ATF6, a regulator of the unfolded protein response. We show that SARS-CoV-2 nsp3.1 directly binds to ATF6 and can suppress the ATF6 stress response. Characterizing the host interactions of nsp3 widens our understanding of how coronaviruses co-opt cellular pathways and presents new avenues for host-targeted antiviral therapeutics.

## INTRODUCTION

Coronaviruses are a family of positive-sense, single-stranded RNA viruses that typically cause upper respiratory infection in humans. Four endemic strains have been characterized that cause symptoms resembling those of the common cold. However, since 2002, three more pathogenic strains have emerged: SARS-CoV in 2002, MERS-CoV in 2012, and SARS-CoV-2, the causative agent of COVID-19, in 2019^1–5^. Some of the differences in pathogenicity can be attributed to differential receptor binding, for example, SARS-CoV and SARS-CoV-2 utilize the angiotensin converting enzyme 2 (ACE2) receptor, while 229E (a common-cold causing strain) uses the human aminopeptidase N receptor^5–7^. At the same time, the engagement of viral proteins with different host proteins or complexes within infected cells is equally critical to understand changes in pathogenicity. These engagements alter the native protein-protein interaction (PPI) architecture of the cell and have been shown to perform various pro-viral functions such as suppression of the type I interferon system for immune evasion purposes^8–10^.

The coronavirus genome is among the largest RNA virus genomes, at approximately 30 kilo base pairs in length. The 3’ third of the genome encodes for the four structural proteins used to construct new virions, as well as several accessory factors shown to be important for pathogenesis. The 5’ two thirds of the genome consists of two open reading frames (orf1a and orf1b) that encode for sixteen non-structural proteins (nsps) that perform a number of functions throughout the viral life cycle, including replication and proofreading of the RNA genome and formation of the replication-transcription complex. The largest of these proteins, at approximately 2000 amino acids, is nsp3. Nsp3 is a large multi-domain protein, of which the papain-like-protease (PL2^Pro^) domain has been most closely studied. In addition to autoproteolysis of the viral polyprotein, the PL2^Pro^ domains possess both deubiquitinase and deISGylation activities^11–13^. Additionally, nsp3 in complex with nsp4 and nsp6 has been shown to be sufficient for formation of the double-membraned vesicles (DMVs) implicated in the CoV replication cycle^14,15^. Expression of the C-terminus of nsp3 and full-length nsp4, while not enough to induce DMV formation, does cause zippering of the ER membrane^16^. However, role(s) of nsp3 outside of the PL2^Pro^ remain less well understood^17^.

Herein, we focused our analysis on four nsp3 homologs from the genus betacoronavirus (hCoV-OC43, MERS-CoV, SARS-CoV, SARS-CoV-2) and one homolog from the genus alphacoronavirus (hCoV-229E). Within the betacoronaviruses, hCoV-OC43 is from clade A, SARS-CoV and SARS-CoV-2 are from clade B, and MERS-CoV is from clade C. The domain organization of nsp3 varies widely among coronavirus genera, and even from strain to strain. Despite the differences, ten regions are conserved across all coronavirus variants: two ubiquitin-like domains (UBLs), a glutamic acid-rich domain, protease domain, two transmembrane regions separated by an ER ectodomain, and two C-terminal Y domains (**Figure S1**).

Affinity purification-mass spectrometry (AP-MS) has been used extensively to characterize the coronavirus interactome, including in two large studies by Gordon et al. to characterize the SARS-CoV-2 host protein interactions and compare these to the SARS-CoV and MERS-CoV interactomes^18,19^. Another study compared the interactomes of isolated SARS-CoV and SARS-CoV-2 PL2^pro^ domains^11^. However, the remaining parts of nsp3 have been missing from SARS-CoV-2 interactome studies thus far, likely because of their complex topology and large size making expression of the protein difficult. To circumvent this problem, we divided the nsp3 protein into three fragments based on earlier interrogation of the SARS intraviral interactome^14,20^. These fragments are referred to as nsp3.1 for the N-terminal fragment, nsp3.2 for the middle fragment, and nsp3.3 for the C-terminal fragment^21^. We expressed each fragment from each of the five viruses listed above (hCoV-229E, hCoV-OC43, MERS-CoV, SARS-CoV, SARS-CoV-2). We employed tandem mass tag (TMTpro 16plex) isobaric tagging technology, which enables highly multiplexed analysis for direct comparison of interactor abundances across homologs from all strains. Previously, we demonstrated the use of AP-MS and TMT technologies in comparing the interactomes of the coronavirus nsp2 and nsp4 proteins from three strains: SARS-CoV-2, SARS-CoV, and hCoV-OC43^22^. We now extend the analysis to the understudied nsp3 protein across additional viral strains. In particular, comparing host protein interactions for homologs from multiple coronavirus strains and genera may provide insight into how the molecular mechanisms of pathogenic coronaviruses differ from endemic ones, as well as the evolution of functions that nsp3 assumes across these strains.

Our study finds that very few interactors are shared among the five strains, although several interactors are common among the mostly conserved SARS variants. Several previously unknown pathways are discovered to be highly enriched with individual variants, such as ERAD processing for SARS-CoV nsp3.2 and nuclear import for 229E nsp3.2. In addition, we find that SARS-CoV-2 nsp3.1 interacts with the unfolded protein response (UPR) transcription factor ATF6 and suppresses the ATF6 pathway in both basal and activated conditions. These discoveries open the door for further work delineating the role of nsp3 in coronavirus infections, with a particular emphasis on the variation in roles between different CoV strains.

## RESULTS

### Expression of nsp3 truncations and AP-MS of CoV nsp3

The orf1a and orf1b open reading frames encode for 16 non-structural proteins which perform crucial roles during the viral life cycle, including replication of the genome and formation of double-membraned vesicles. Non-structural protein 3 (nsp3) is the largest of the 16 non-structural proteins; the full-length protein is 1586 – 1945 amino acids long (177-217 kDa) (**Figure 1A**). While the domain organization is different among CoV variants, several domains are shared, including multiple ubiquitin-like-domains, at least one highly conserved papain-like protease (PL2^Pro^) domain, and an ER luminal domain postulated to be important for nsp4 binding and double membrane vesicle (DMV) formation (**Figure 1B-C, Figure S1**).

**Figure 1.**
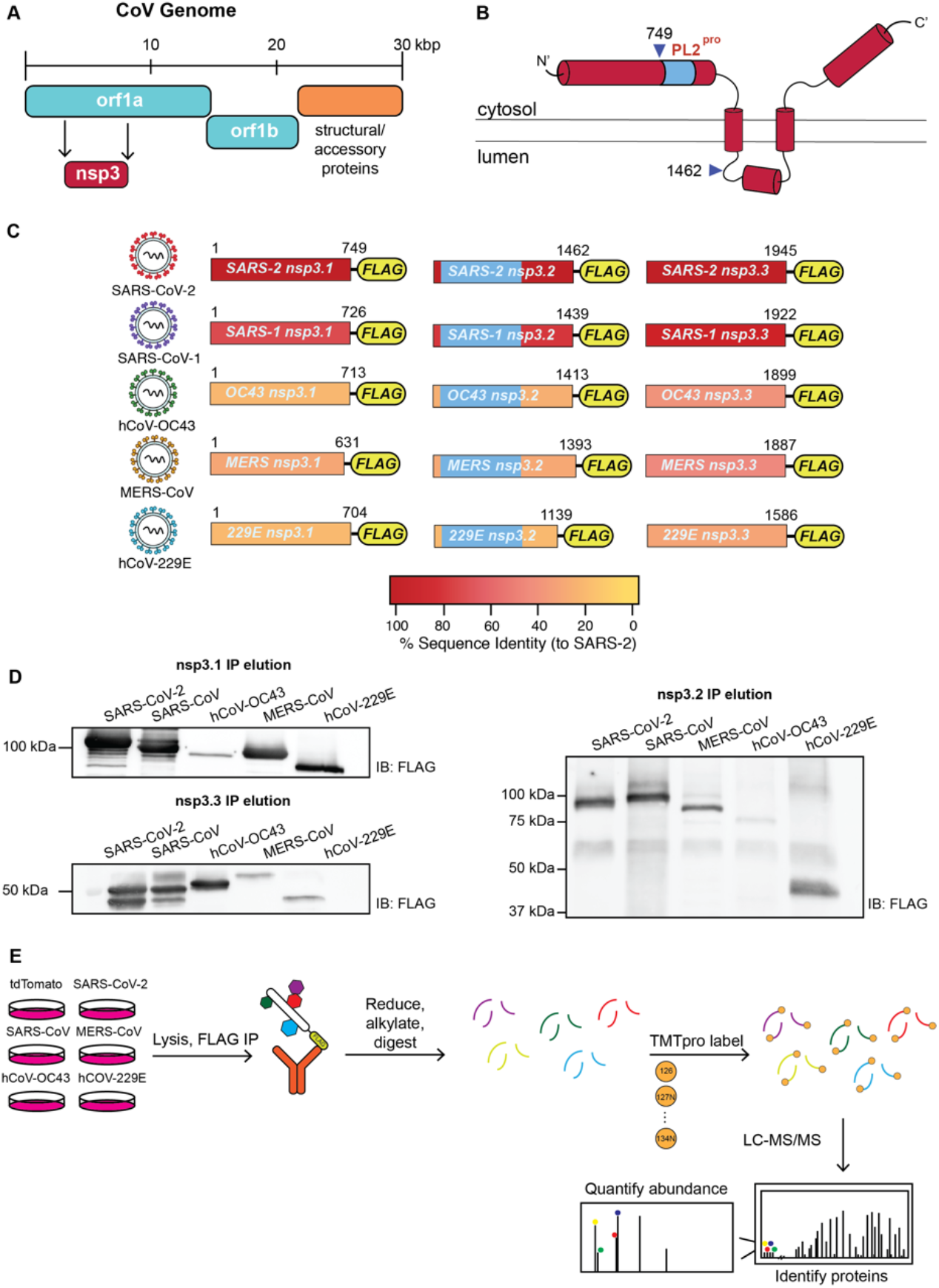
Design and expression of CoV nsp3 truncations for affinity-purification mass spectrometry (AP-MS). A) Coronavirus (CoV) genome schematic indicating the regions encoding orf1a, orf1ab, and structural/accessory proteins. Nsp3 is encoded within orf1a. B) General schematic of SARS-CoV-2 nsp3 protein topology. Nsp3 has two single-span transmembrane regions anchoring the protein in the ER membrane, with a small luminal domain and both N and C terminal regions in the cytosol. The conserved papain-like protease (PL2^pro^) domain is on the N-terminal cytosolic portion. For this study, the nsp3 protein was truncated into three fragments: nsp3.1 (1-749), nsp3.2 (750-1462), and nsp3.3 (1463-1945), numbering corresponding to SARS-CoV-2 nsp3. C) The nsp3 truncations for homologs from all five human coronaviruses used in this study. All fragments contain a C-terminal FLAG tag for affinity-purification. Percent sequence identity compared to SARS-CoV-2 is indicated. The PL2^pro^ domain in nsp3.2 homologs is highlighted in light blue. D) Western blotting of immunopurified nsp3.1, nsp3.2, and nsp3.3 homologs after transient transfection in HEK293T cells. E) AP-MS workflow to identify virus-host protein interactions of nsp3 fragment homologs. HEK293T cells were transfected with corresponding homologs and lysates were immunopurified using anti-FLAG beads to enrich for viral proteins in complex with host interactors. Proteins were reduced, alkylated, and digested with trypsin. Peptides were then labeled with tandem mass tags (TMTpro) and analyzed by tandem mass spectrometry (LC-MS/MS) to both identify and quantify host interactors.

Initial attempts at expression of the full-length construct for SARS-CoV-2 nsp3 were unsuccessful, likely due to the complex structure and topology of the full-length protein. To study the interactome, we thus divided the protein into three portions based on a prior interactome study carried out on similar truncations of the SARS-CoV nsp3 homolog^20^. The N-terminal fragment (nsp3.1), comprising the first ubiquitin-like-domain through the SARS unique domain, is expected to localize exclusively to the cytosol (**Figure 1C**, **Figure S1**). The second portion (nsp3.2), starting before the second ubiquitin-like domain and ending just after the first transmembrane region, includes the papain-like-protease (PL2^Pro^) domain. The C-terminal fragment (nsp3.3) starts with the ER-localized 3Ecto-domain, includes the second transmembrane region and ends with the C-terminus of the protein. Based on multiple sequence alignments, similar N-terminal, middle, and C-terminal constructs were created for the SARS-CoV, MERS-CoV, hCoV-OC43, and hCoV-229E nsp3 homologs (**Figure S1**). Each construct contains a C-terminal FLAG tag for affinity purification (**Figure 1C**).

Constructs were transfected into HEK293T cells, and expression was confirmed by immunoprecipitation and immunoblotting for the FLAG tag and/or detection by mass spectrometry. Although kidney cells do not represent a primary tissue target of the virus, previous work has identified HEK293T cells as appropriate cell lines to recapitulate relevant CoV protein interactions with host factors^18,22^. While the constructs showed variable expression levels, all fragments were reproducibly detectable by mass spectrometry, and peptides coverage spanned the length of each fragment (**Figure 1D**). For native co-immunoprecipitations with host interactors, protein constructs were expressed, lysed in mild detergent buffer to maintain interactions, and co-immunoprecipitated from lysates using anti-FLAG beads. After confirmation of IP by silver stain, samples were reduced, alkylated, and trypsin digested. Samples were labeled using TMTpro 16plex reagents for MS2 quantification of peptides abundances^23^ (**Figure 1E**). Between 2-4 co-immunoprecipitation replicates for each construct were pooled into a single TMTpro 16plex run, along with mock IPs from tdTomato transfected cells to establish the background signal. The final datasets consisted of 184 IPs across the 16 constructs (tdTomato & 15 viral protein constructs) (**Figure S2**).

After identification and quantification of interactors using Proteome Discoverer, a variable cutoff method was used to determine high- and medium- confidence interactors for each construct based on enrichments compared to the tdTomato background control.

### Comparison of CoV nsp3.1-host interactors

The N-terminal fragments of nsp3 contain a conserved ubiquitin-like domain (Ubl1), a highly variable or acidic-rich region, a conserved macrodomain (Mac1), followed by strain-specific domains (**Figure 2A, Figure S1**). In hCoV-229E and hCoV-OC43, this fragment includes a papain-like protease domain (termed PL1^Pro^). This first PL1^Pro^ domain is absent from all viruses in clades B and C of the betacoronaviruses, hence its absence in MERS-CoV and the two SARS variants. In contrast, SARS strains contain a SARS-unique domain (SUD) consisting of 3 sequential macrodomains. Overall, this region of nsp3 is most variable from strain to strain, despite several of the predicted domains being conserved (**Figure 1B-C, Figure S1**).

**Figure 2.**
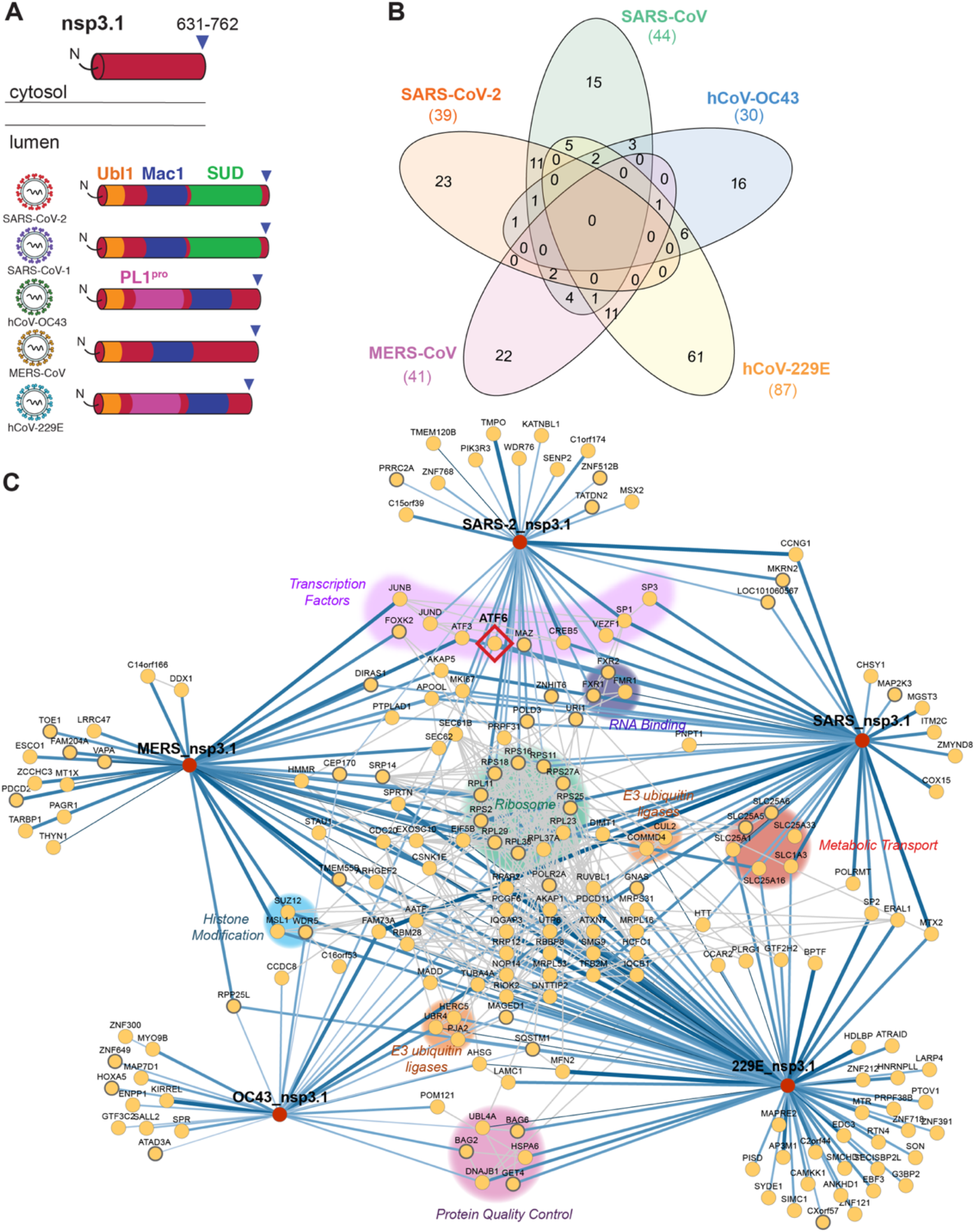
Identification of CoV nsp3.1 host interactors. A) Schematic of nsp3.1 topology for all five CoV homologs. Nsp3.1 is a cytosolic fragment, comprising residues from 1 to 631-762. All fragment homologs contain a ubiquitin-like domain (Ubl1, yellow) and a conserved macrodomain (Mac1, blue). SARS-CoV-2 and SARS-CoV-1 contain a SARS-unique domain (SUD, green), while hCoV-OC43 and hCoV-229E contain a papain-like protease domain (PL1^pro^, pink). B) Venn diagram showing the number of unique and shared host interactors amongst all five CoV nsp3.1 homologs. Total interactors for each homolog are shown in parentheses. C) Network plot of virus-host interactors. Individual nsp3.1 homologs are shown as red circles, while host interactors are shown as yellow circles. Blue lines indicate virus-host protein interactions, where line width and shade are wider/darker for more highly enriched interactions. Grey lines indicate known host-host protein interactions from the STRING database. Notable clusters of host proteins are highlighted. The transcription factor ATF6 (red diamond) interacts with SARS-CoV-2, SARS-CoV-1, and hCoV-OC43 and was subjected to later functional follow-up (**Figure 5**).

The final dataset for the N-terminal fragment analysis combined 5 mass spectrometry runs containing 62 co-IP samples (tdTomato 15, SARS-CoV-2 13, SARS-CoV 9, OC43 9, MERS 8, 229E 8) (**Figure S2A**). We identified a robust set of high-confidence interactors for each fragment: 39 for SARS-CoV-2, 44 for SARS-CoV, 41 for MERS-CoV, 30 for hCoV-OC43, and 87 for hCoV-229E (**Figure S3**). Of these high-confidence interactors, none were common across all 5 strains, and none were common to all betacoronaviruses (**Figure 2B**). Fourteen interactors were common between the two SARS variants, and 2 of these 14 interactors, MKI67 and ATF3, were also shared with the MERS fragment. MKI67 is required to maintain individual chromosomes in the cytoplasm during mitosis, and ATF3 is a cyclic AMP-dependent transcription factor which negatively regulates the cellular antiviral response. Lastly, we identified 25 high confidence interactors unique to SARS-CoV-2, 16 unique to SARS-CoV, 17 unique to hCoV-OC43, 62 unique to hCoV-229E, and 23 unique to MERS-CoV. Notably contained in this fragment is the SARS-unique domain, which is exclusively found in SARS-CoV and SARS-CoV-2 nsp3. As noted, 14 proteins were identified as interactors of both fragments. Eleven of these interactors were exclusive to the SARS variants, including several ribosomal proteins (RPL38, RPS11, RPS2, RPS16) and assorted factors (FMR1, MKRN2, SP1, VEZF1, CREB5, CCNG1, and PEX11B).

We also identify 9 shared interactors between OC43 and 229E, both of which have a PL1^pro^ domain (UBR4, TUBA4A, SQSTM1, RPP25L, MADD, DNAJB1, BAG6, AHSG, AATF). BAG6 and DNAJB1 are co-chaperones components involved in protein quality control ^24,25^, while UBR4 is an E3 ubiquitin ligase involved in membrane morphogenesis and SQSTM1 is an autophagy receptor^26,27^. These shared interactors may represent possible targets of the PL1^pro^ domains in OC43 and 229E.

As an alternative method to map shared and unique interactors for the different strains, we performed hierarchical clustering of the grouped protein abundance Z-scores for all identified high-confidence interactors (**Figure S4**). Consistent with the Venn diagram, we observed a cluster of shared interactors for the SARS construct (cluster 4), distinct interactors of 229 (cluster 5) and MERS (cluster 3). A large cluster (7) contained interactors present in four of the strains but absent in SARS-CoV-2.

To ascertain how these proteins cluster into cellular pathways with potential pro- or anti-viral roles, we grouped the high-confidence interactors into a network plot. To highlight connections between the binding partners, known protein interactions based on the STRING database were included (**Figure 2C**). Several groups of associated interactors emerged, including a large cluster of eleven ribosomal proteins (RPS2, RPS11, RPS16, RPS18, RPS25, RPS27A, RPL11, RPL23, RPL29, RPL37A, RPL38). Other notable clusters include factors involved in protein quality control (BAG2, BAG6, HSPA6, DNAJB1, UBL4A, GET4), metabolic transport (SLC1A3, SLC25A1, SLC25A5, SLC25A6, SLC25A16, SLC25A33), histone modification (SUZ12, MSL1, WDR5), RNA-binding proteins (FXR1/2, FMR1), and E3 ubiquitin ligases (CUL2, COMMD4, HERC5, UBR4, PJA2). Additionally, ten transcription factors were identified with at least one interaction to severely pathogenic betacoronaviruses. These include VEZF1, a transcription factor for IL-3^28^, ATF3, a broad negative regulator of NF-κB, IL-4, IL-5, IL-13 expression^29^, and ATF6, which turns on expression of the ATF6 branch of the unfolded protein response (UPR)^30^.

As a last method to identified pathways represented by the interactors, we filtered the list of proteins through EnrichR to determine the most common gene ontology terms related to biological processes among each interactome (**Figure S5**). For both SARS variants, as well as MERS-CoV, we observed a strong enrichment for several ribosomal-related processes, including SRP-dependent cotranslational protein targeting, ribosome biogenesis, and rRNA metabolic processes. These processes were not observed in the less pathogenic hCoV-OC43 and hCoV-229E strains. For SARS-CoV, but not SARS-CoV-2, we also observed several mitochondrial related processes, such as mitochondrial transport, and mitochondrial RNA metabolic processes. hCoV-OC43 showed enrichment of factors associated with an upregulation of transcription from RNA polymerase II factors. Lastly, despite having the highest overall number of high confidence interactors, no individual pathways showed enrichment at the high confidence interactor level for hCoV-229E.

### Comparison of nsp3.2 CoV interactors

Next, we turned to the middle fragment of the nsp3 constructs, which includes the highly conserved PL2^Pro^ domain, the protease responsible for the self-cleavage of several coronavirus non-structural proteins from the orf1a/b polypeptide^31^. This domain also has deubiquitination and de-ISGylation activity, an activity well-studied in the pathogenic coronaviruses, but less studied in the endemic strains^32,33^. The importance of this domain for both viral polypeptide processing and remodeling of host ubiquitination/ISGylation modifications make it an intriguing target for drug development. Also included in this fragment are the second ubiquitin-like domain UBL2, the first transmembrane domain, and the first portion of the ER ectodomain (**Figure 3A**).

**Figure 3.**
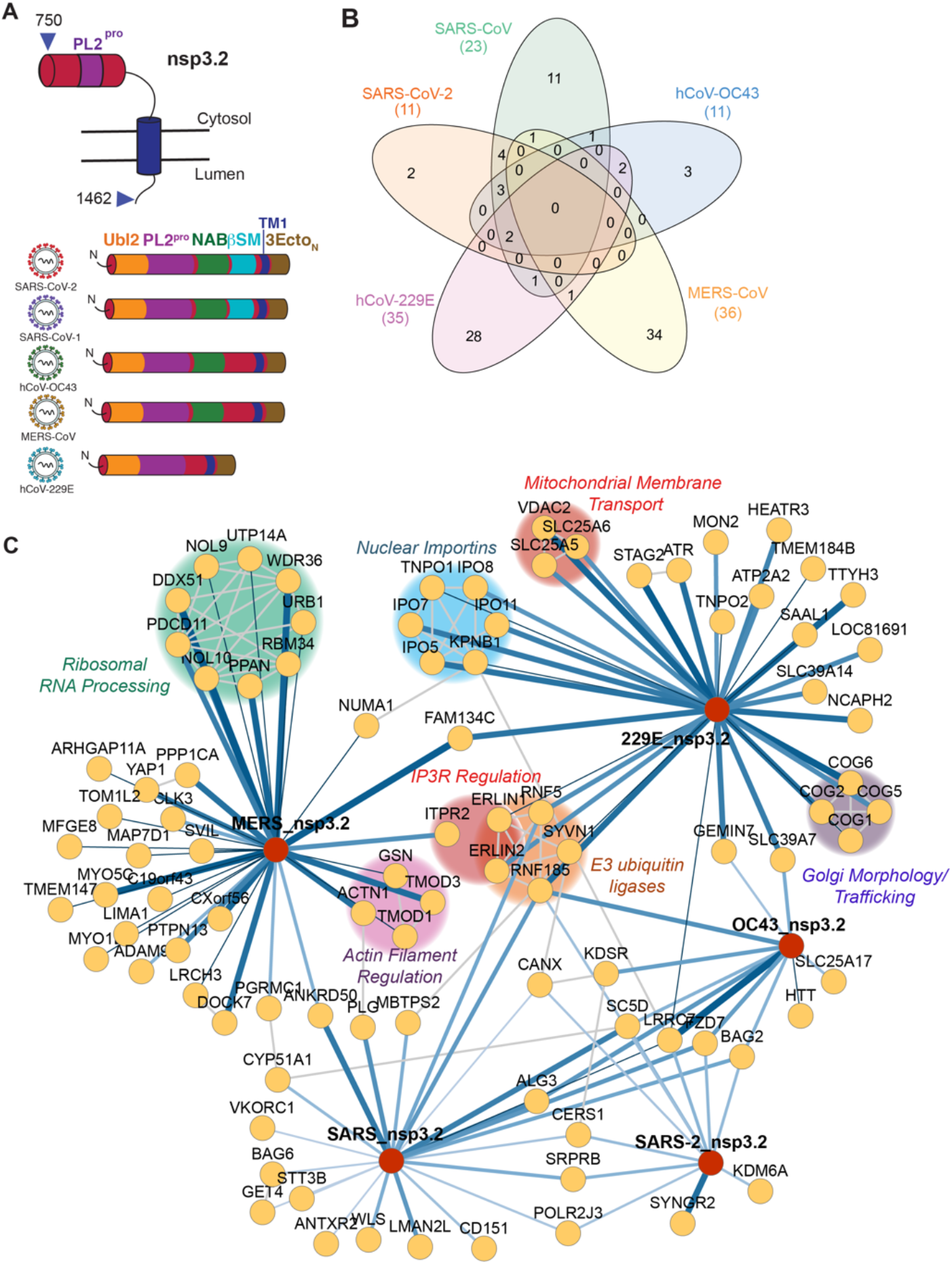
Comparative interactomics of CoV nsp3.2 homologs. A) Schematic of nsp3.2 protein topology for all five CoV homologs. Nsp3.2 contains both cytosolic, transmembrane, and luminal regions. All homologs contain a ubiquitin-like domain (Ubl2, yellow), a papain-like protease domain (PL2^pro^, purple), a transmembrane region (TM1, blue), and the N-terminal portion of the ectodomain (3Ecto_N_, brown). Betacoronavirus homologs also contain a nucleic acid binding domain (NAB, green), while SARS strains also have a betacoronavirus-specific marker (βSM, light blue). B) Venn diagram showing the number of unique and shared host interactors amongst all five CoV nsp3.2 homologs. Total interactors for each homolog are shown in parentheses. C) Network plot of virus-host interactors. Individual nsp3.2 homologs are shown as red circles, while host interactors are shown as yellow circles. Blue lines indicate virus-host protein interactions, where line width and shade are wider/darker for more highly enriched interactions. Grey lines indicate known host-host protein interactions from the STRING database. Notable clusters of host proteins are highlighted.

The final dataset for the middle fragment analysis combined 3 mass spectrometry runs containing 59 co-IP samples (tdTomato 10, SARS-CoV-2 11, SARS-CoV 11, hCoV-OC43 10, MERS-CoV 10, hCoV-229E 7) (**Figure S2B**). In total, we identified 11 high confidence interactors for SARS-CoV-2, 23 for SARS-CoV, 11 for hCoV-OC43, 36 for MERS-CoV, and 35 for hCoV-229E (**Figure 3B, Figure S6**). Nine proteins were observed as high confidence interactors for both SARS-CoV and SARS-CoV-2. No proteins were observed to be common among all 5 strains at the high confidence level (6 at the medium confidence level), and no proteins were observed to be exclusive interactors of betacoronaviruses at the high confidence level (2 at the medium confidence level). Lastly, we identified 2 high confidence interactors unique to SARS-CoV-2, 11 unique to SARS-CoV, 3 unique to OC-43, 28 unique to 229E, and 34 unique to MERS.

We carried out hierarchical clustering of the grouped abundance Z-scores for all high-confidence interactors. The resulting heatmap (**Figure S7**) more clearly highlights a cluster of shared interactions for all five strains (cluster 2), although intensities vary, which could explain why not all may have passed our stringent high-confidence cutoff. Cluster 1 contains binding partners found in four of the strains, but absent in MERS. A striking observation were two large clusters of strain-specific interactors for 229E (cluster 3) and MERS (cluster 4) respectively.

We also grouped the high-confidence interactions into a network plot highlighting previously known interactions from String DB (**Figure 3C**). We identify several ER-associated degradation (ERAD) components as high confidence interactors of SARS-CoV nsp3.2. While some of these components were identified as high confidence interactors of SARS-CoV-2 as well, the number and magnitude were both much less. Analysis of GO terms associated with both the medium and high confidence interactors confirmed the enrichment of the ERAD machinery, as well as several related processes such as membrane proteolysis and cellular response to ER stress (**Figure S8**). The 229E fragment revealed a high number of unique interactors related to nuclear importins and translocation of proteins into the nucleus (IPO5, IPO7, IPO11, IPO8, KPNB1, TNPO1). A recent report identified a role of MHV nsp3 in forming pores across the DMV membrane in coronavirus infection, potentially for the purpose of dsRNA export from the DMVs^34^. The nuclear importins could be additional host factors co-opted by 229E or alphacoronaviruses specifically for a similar purpose. Additional interactors of the 229E fragment include ERLIN1 and ERLIN2, which we previously identified as being interactors of both SARS-CoV-2 and SARS-CoV nsp2 and nsp4. These proteins are also associated with ERAD, most notably in regulating ubiquitination and degradation of the inositol 1,4,5-triphosphate receptor IP3R^35^. Lastly, other unique interactor of hCoV-229E nsp3.2 included a cluster of mitochondrial membrane transporters (SLC25A6, SLC25A5, VDAC2), as well as a cluster of subunits of the conserved oligomeric Golgi (COG) complex involved in intra-Golgi mediated vesicle transport. COG6 was identified in a recent CRISPR screen as essential for hCoV-229E replication^36^, and the intra-Golgi mediated vesicle transport was a pathway enriched only in our hCoV-229E nsp3.2 dataset, not appearing for the other strains (**Figure S8**).

Lastly, for MERS nsp3.2. we identified a complex of ribosomal RNA (rRNA) processing factors as highly unique interactors, including DDX51, NOL9, NOL10, UTP14A, WDR36, URB1, RBM34, PPAN, and PDCD11 (**Figure 3C**). Consistent with this finding, RNA processing and ribosome biogenesis were the most highly-enrichment GO terms specific to the MERS nsp3.2 fragment (**Figure S8**). Other viruses have been shown to target ribosomal biogenesis to facilitate infection, such as Human cytomegalovirus (HCMV) and HIV-1^37,38^. The specificity of these interactors for MERS nsp3.2 is striking and may represent a unique replication strategy to modulate host protein synthesis.

### Comparison of nsp3.3 CoV interactors

The C-terminal fragments begin with the second half of the ER ectodomain, continuing through the second transmembrane region and Y domains, ending with the nsp3/nsp4 cleavage site. The Y domains, although one of the more conserved domains across the 5 strains, remains largely unstudied. In co-expression experiments of individual SARS-CoV-2 nsp3 fragments and nsp4, we observed co-immunoprecipitation of this C-terminal fragment with nsp4 (**Figure S9**). In contrast, the N-terminal and middle fragments did not co-immunoprecipitate with nsp4.

The final dataset for the middle fragment analysis combined 5 mass spectrometry runs containing 63 co-IP samples (tdTomato 15, SARS-CoV-2 14, SARS-CoV 9, hCoV-OC43 9, MERS-CoV 8, hCoV-229E 8) (**Figure S2C**). In total, we identified 75 high confidence interactors for SARS-CoV-2, 66 for SARS-CoV, 52 for hCoV-OC43, 49 for MERS-CoV, and 76 for hCoV-229E (**Figure 4B**, **Figure S10**). One protein, SLC39A6, a zinc transporter, was observed as a high confidence interactor for all five strains (68 proteins observed as common medium confidence interactors). Five proteins, DERL3, TMEM33, SC5D, CERS1, and ALG8, were observed as high confidence interactors unique to the betacoronavirus strains. DERL3 is a functional component of the ERAD system, TMEM33 is involved in tubular ER network organization as well as being a component of the IRE1 and PERK stress response pathways, SC5D is involved in cholesterol biosynthesis, CERS1 is involved in lipid biosynthesis, and ALG8 is involved in N-glycan biosynthesis. Lastly, we identify 48 high confidence interactors unique to SARS-CoV-2, 17 unique to SARS-CoV, 20 unique to hCoV-OC43, 41 unique to hCoV-229E, and 18 unique to MERS-CoV (**Figure 4B**).

**Figure 4.**
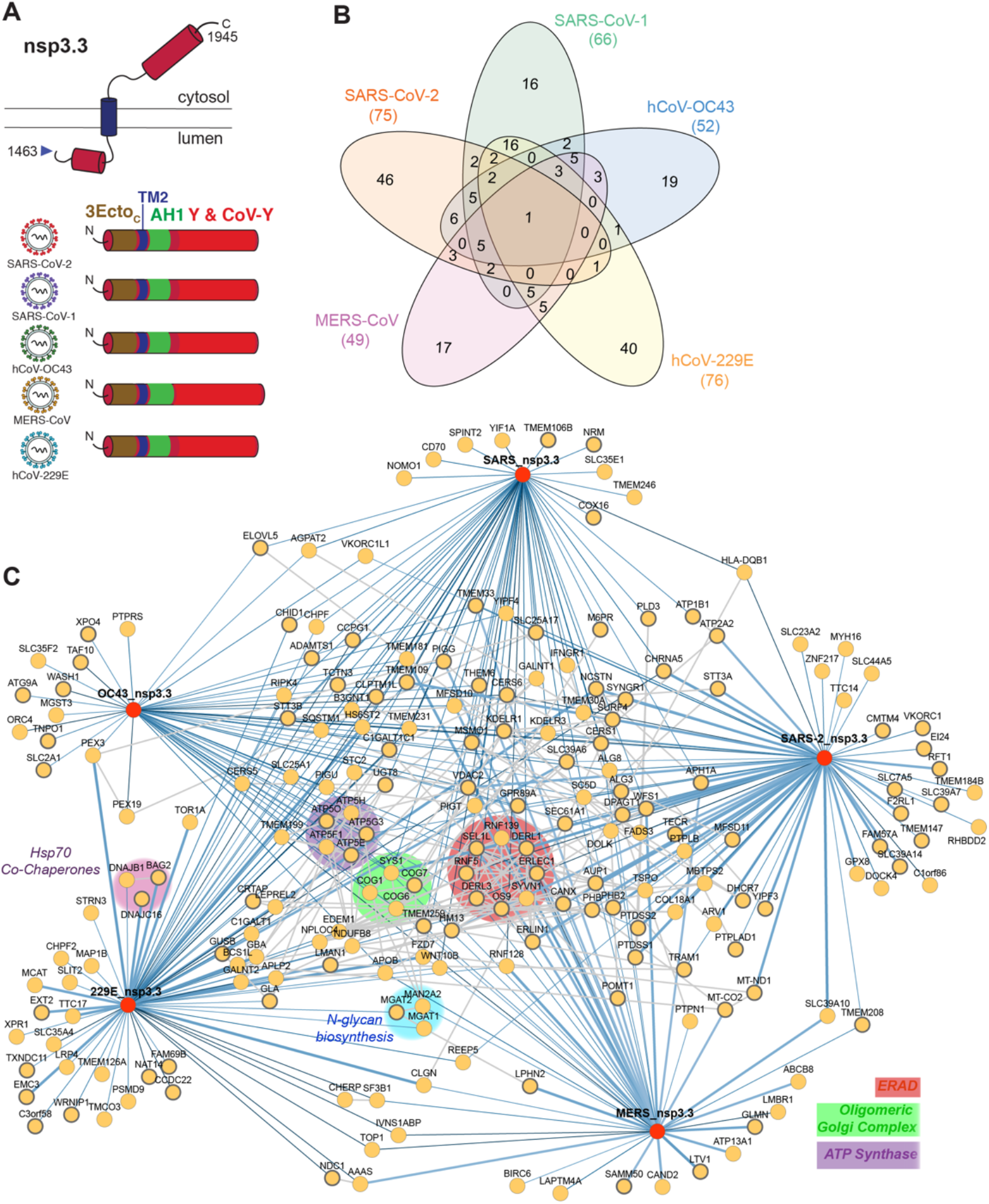
Identification of CoV nsp3.3 host interactors. A) Schematic of nsp3.3 protein topology for all five CoV homologs. Nsp3.3 contains both cytosolic, transmembrane, and luminal regions. All homologs contain the C-terminal portion of the ectodomain (3Ecto_C_, brown), the second transmembrane region (TM2, blue), a likely amphipathic helix (AH1, green), and a Y&CoV-Y domain (red). B) Venn diagram showing the number of unique and shared host interactors amongst all five CoV nsp3.3 homologs. Total interactors for each homolog are shown in parentheses. C) Network plot of virus-host interactors. Individual nsp3.3 homologs ar e shown as red circles, while host interactors are shown as yellow circles. Blue lines indicate virus-host protein interactions, where line width and shade are wider/darker for more highly enriched interactions. Grey lines indicate known host-host protein interactions from the STRING database. Notable clusters of host proteins are highlighted.

We carried out hierarchical clustering of the grouped abundance Z-scores for all high-confidence interactors. The resulting heatmap (**Figure S11**) highlights 6 clusters of interactors, including a cluster of shared interactors for all five strains (cluster 4), a cluster of shared betacoronavirus interactors (cluster 5), and a cluster of shared SARS interactors (cluster 6).

Despite these C-termini being the most conserved of the fragments across strains, there was a large divergence observed in the enriched pathways when searched for enriched GO-terms (**Figure S12**). In all strains except hCoV-OC43, ERAD was a highly enriched pathway. It has been postulated that nsp3, in tandem with nsp4 and nsp6, hijacks ERAD-tuning vesicles during double-membrane vesicle (DMV) formation, after ERAD factors EDEM1 and OS-9 were shown to co-localize with MHV-derived DMVs in infected cells^14,39^. Our results may provide insight into a role for nsp3 in modulating this machinery.

Unexpectedly, the biological process most enriched for the hCoV-229E fragment was protein N-linked glycosylation. N-linked glycosylation sites are localized at the N-terminus of the 3.3 fragment in the ectodomain. One of these interactors, MGAT1 (unique to the hCoV-229E interactome), was identified in the CRISPR screen as being an essential gene for hCoV-229E infection^36^. In addition, we identify STT3B, a catalytic component of the oligosaccharyl transferase complex, as a high confidence hCoV-229E nsp3.3 interactor.

The patterns observed in our GO term analysis were consistent with network plots of high-confidence interactors (**Figure 4C**). Multiple homologs shared interactors involved in ERAD (RNF139, DERL1, DERL3, ERLEC1, SYNV1, OS9, RNF5, SEL1L) and the oligomeric Golgi complex (COG1, COG6, COG7, SYS1). A cluster of five components of the mitochondrial ATP synthase complex (ATP5E, ATP5G3, ATP5H, ATP5F1, and ATP5O) was enriched for hCoV-OC43 nsp3.3, while clusters of Hsp70 co-chaperones (DNAJB1, DNAJC16, BAG2) and *N*-glycan biosynthesis factors (MGAT1, MGAT2, MAN2A2) were enriched for hCoV-229E nsp3.3. Interestingly, a large swath of identified high-confidence interactors are interconnected based on STRING analysis but do not cluster into distinct biological categories outside of those previously mentioned.

### SARS-CoV-2 nsp3.1 interacts with ATF6 and suppresses the ATF6 branch of the Unfolded Protein Response (UPR)

An intriguing observation was the shared interaction of the nsp3.1 fragment from SARS-CoV-2, SARS-CoV and hCoV-OC43 with ATF6 (**Figure 2C**), a transmembrane protein located across the ER membrane serving as one of the sensors of the ER unfolded protein response (UPR). Upon activation by ER stress, ATF6 is trafficked to the Golgi apparatus, where site-1 and site-2 proteases (SP1 & MBTPS2) cleave the protein, releasing an active transcription factor that serves to upregulate chaperones and other proteostasis factors which aid in relieving ER stress^30^. Intriguingly, we also observed MBTPS2 as an interactor of SARS-CoV nsp3.2. MSTPS2 was found in a recent CRISPR to be an essential gene in the replication of SARS-CoV-2^36^. Coronaviruses are known to upregulate the UPR. Specifically, the SARS-CoV spike protein was shown to be sufficient to upregulate transcription of factors such as GRP78/BiP and GRP94^40,41^, both known to be downstream targets of ATF6^42,43^. Furthermore, SARS-1 Orf8 can activate ATF6 and promote UPR induction^44^, but a role for nsp3 in UPR modulation during virus infection is not known.

As functional validation of the interaction of the N-terminal fragment of SARS-CoV-2 nsp3 with ATF6, we used a HEK293T cell line expressing a doxycycline-inducible 3xFLAG-ATF6 construct^45^. The affinity tag of the nsp3 N-terminal construct was replaced with a 2x strep tag (2xST) to allow for complementary immunopurifications (IP) and detection. We transiently transfected 3xFLAG-ATF6 cells with SARS-CoV-2 nsp3.1-2xST and carried out reciprocal co-IPs to validate the interactions (**Figure 5A**). We detected some background nsp3.1-2xST protein in the control (DMSO-treated) FLAG-IP where ATF6 was not induced. This was likely due to some leaked expression. However, the viral protein was noticeably enriched when 3xFLAG-ATF6 was robustly induced with doxycycline. ATF6 was also observed in reciprocal Co-IPs (**Figure 5A**), further validating the interaction between SARS-CoV-2 nsp3.1 and ATF6.

**Figure 5.**
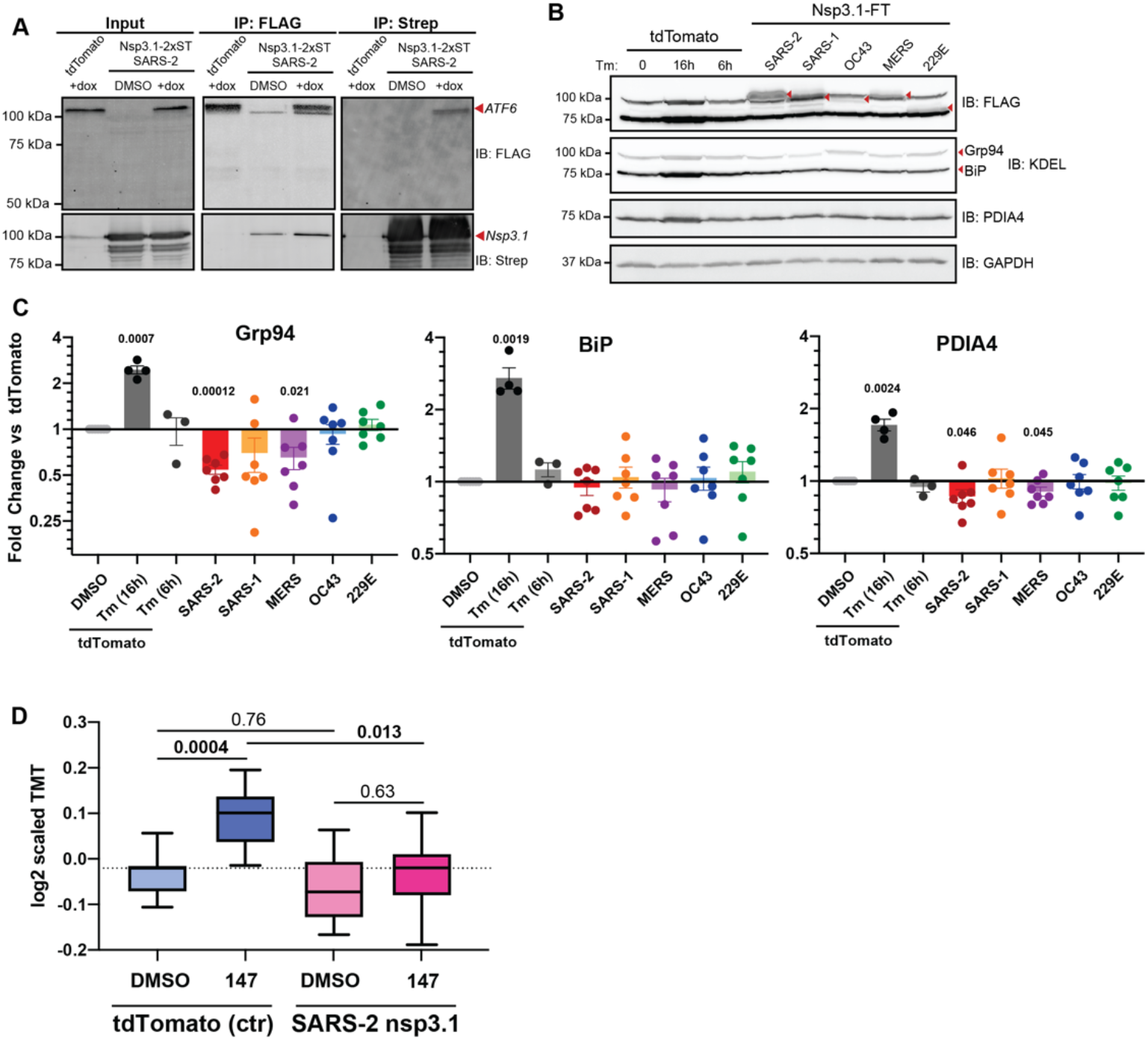
SARS-CoV-2 nsp3.1 interacts with ATF6 and suppresses the ATF6 branch of the Unfolded Protein Response (UPR). A) Representative co-immunopurification (IP) western blots of 3xFT-ATF6 cells transfected with SARS-CoV-2 nsp3.1-2xST or tdTomato (control), lysed, and immunopurified for either FLAG or 2xStrepTag. Cells were treated with 100 nM doxycycline to induce 3xFT-ATF6 expression 24 h pre-harvest. Input and IP elution blots were probed with both anti-FLAG and anti-StrepTag antibodies. *n* = 3 B) Representative western blots of HEK293T cells transfected with nsp3.1-FT homologs or tdTomato (control), lysed, and probed for FLAG, ATF6 branch markers (Grp94, BiP, PDIA4), and GAPDH as a loading control. Control cells were either treated with DMSO or 6 mg/mL Tunicamycin (Tm) at 6 or 16 h pre-harvest to induce an ER stress response. TdTomato 16 h pre-harvest, *n* = 4; TdTomato 6 h pre-harvest, *n* = 3; all others, *n* = 7. C) Quantification of western blots shown in (B). Error bars indicate average ±SEM. Paired student T-tests were used to test for significance between samples and tdTomato+DMSO control, with *p*-values < 0.05 shown. D) Box-and-whisker plots of ATF6-regulated protein abundance measured by quantitative proteomics. HEK293T cells were transfected with tdTomato (control) or SARS-CoV-2 nsp3.1-FT and treated with DMSO or 10 μM **147** for 16 h pre-harvest. Shown are the distribution of scaled log2 TMT intensities for ATF6-regulated proteins based on published genesets^43^. A one-way ANOVA with Geisser-Greenhouse correction and post-hoc Tukey’s multiple comparison test was used to determine significance. Adjusted *p*-values are shown. *n* = 3 biological replicates in a single mass spectrometry run.

Given this observation of SARS-CoV-2 nsp3.1 directly binding to ATF6, we sought to determine if this interaction could constitute a mechanism by which the UPR may be modulated. Cells were transfected with all five nsp3.1 homologs and immunoblotting was performed for known ATF6-specific target genes (GRP78/BiP, GRP94, PDIA4) to determine if upregulation of these proteins occurred. The nsp3.1 fragments were robustly expressed, but no notable upregulation of GRP78/BiP, GRP94 or PDIA4 was observed (**Figure 5B-C**). Instead, we saw a small but significant reduction in protein levels for GRP94 and PDIA4 with SARS-CoV-2 and MERS-CoV nsp3.1. The same experiment was performed to monitor changes of RNA transcript level, but similarly no upregulation of *BiP*, *PDIA4* or *ERDJ4* (IRE1 marker) transcripts was seen (**Figure S13**). We also did not observe any site-1/2 protease mediated-cleavage of ATF6 in SARS-CoV-2 nsp3.1-transfected samples (**Figure 5A**).

Given the small reduction in GRP94 and PDIA4 protein levels (**Figure 5B-C**), an alternative possibility was that the nsp3.1 binding event could suppress ATF6 activation. To further test this possibility, cells were transfected with the respective nsp3.1 constructs and treated with compound **147**, a small molecule ER proteostasis regulator that preferentially activates the ATF6 branch of the UPR^46,47^. Importantly, **147** treatment mimics the smaller degree of UPR activation observed during CoV infection compared to treatment with global ER stressors, such as tunicamycin or thapsigargin^40,48^. We profiled global expression changes using quantitative proteomics to gain a broader insight into the regulation of UPR target genes based on larger genesets of ATF6-regulated targets, as well as targets of the other UPR branches (IRE1/XBP1s and PERK)^43^. SARS-CoV-2 nsp3.1 resulted in a slight downregulation of ATF6 targets compared to a control transfection, but this was not significant (**Figure 5D**). We observed robust activation of ATF6 target genes when control-transfected cells were treated with **147**. In contrast, no significant upregulation of ATF6 targets was observed in the nsp3.1 transfected cells treated with **147**, confirming the suppression of the ATF6 response (**Figure 5D**). To ensure that the suppression was specific to ATF6 targets, we also analyzed the response of **147** treatment and SARS-CoV-2 nsp3.1 expression on other UPR branches and the cytosolic heat shock response (**Figure S14**). We did not observe any change in expression in response to **147** or nsp3.1 further confirming that the regulation is specific to ATF6.

## DISCUSSION

As evidenced by its many domains, nsp3 likely serves a multitude of roles within the coronavirus replication cycle. Some of these roles are well characterized, such as the requirement of nsp3 for formation of double-membraned vesicles and the papain-like protease function in autocleavage of the orf1a polypeptide^14–17,49,50^. Other roles remain less defined, and the interactome of the individual nsp3 fragments contained herein may serve to help delineate the role of this protein across several CoV strains. Using tandem mass tags (TMTpro 16plex), we are able to directly identify and compare the abundance of interactors across five coronavirus strains, including two different genera and three different betacoronavirus clades. Given the frequent emergence of severely pathogenic coronavirus strains over the past 20 years, it is becoming increasingly apparent that a deeper knowledge of the molecular mechanisms of coronavirus replication is needed, to understand how we may better prepare therapeutics for potential future strains of these viruses.

The N-terminal portion of the protein (nsp3.1) is the only fragment lacking transmembrane domains and exclusively localizes to the cytosol. This could explain the high expression levels seen with this fragment, although there was still some variation among the expression efficiency for the homologs from different viral strains. An interesting divergence observed in the nsp3.1 dataset is the enrichment of pathways related to mitochondrial transport and metabolism in the SARS-CoV dataset that is absent in the SARS-CoV-2 dataset. Given the two fragments share high sequence identity, it is noteworthy when one pathway is so highly enriched in one fragment versus the other, pointing to rapid divergent evolution. Other SARS-CoV-2 proteins have been shown to interact with several components of the mitochondria^18,19^, including our earlier nsp2/4 dataset that found both SARS-CoV and SARS-CoV-2 proteins interacting with mitochondria-associated membrane factors involved in controlling calcium flux between ER and mitochondria^22^. Interestingly, the most highly enriched pathways in both of the SARS variants are also enriched (to a similar magnitude) in the MERS-CoV nsp3.1 fragment pointing towards conserved functions among the pathogenic variants.

The C-terminal portion of the nsp3.1 fragment for SARS-CoV and SARS-CoV-2 consists of the SARS unique domain (SUD). While a definitive function of this domain has not been established in the coronavirus replication cycle, prior studies described a role for macrodomains within the SUD binding G-quadruplexes, strings of RNA containing multiple guanosines^49,51,52^. While no RNA-binding pathways were observed as enriched with either of the SARS nsp3.1 fragments, potentially related processes such as mRNA catabolism, rRNA metabolism, and ribosome biogenesis were significantly enriched. This pathway was also observed in the MERS-CoV nsp3.1 interactor set, potentially pointing to a broader role for pathogenic strains. The singular pathway shown to be enriched only in the two SARS variants was “positive upregulation of transcription in response to ER stress”, which prompted us to investigated the interactions and regulation of this fragment with the UPR sensor ATF6, which is discussed further below.

The middle fragment yielded the smallest number of overall interactors of the three fragments, across all five strains. Generally, expression levels of this fragment were also lower than the termini, which may have limited detection of interactors. However, all fragments and interactors were still reproducibly detectable by mass spectrometry. Surprisingly, we observed highly distinct pathway enrichment from the MERS-CoV and hCoV-229E fragments showing substantial interactions with ribosomal RNA processing proteins and nuclear import proteins, respectively. Interactions of other coronavirus proteins with nuclear transport machinery have been observed, for instance nsp9 with the nuclear pore complex and nsp1 with mRNA export machinery^18,53^. Orf6 of SARS-CoV and SARS-CoV-2 inhibits STAT1 signaling by blocking nuclear import of phosphorylated STAT1. Nuclear localization of SARS-CoV and SARS-CoV-2 orf3b has been shown, as well as for N proteins^54–56^. In contrast, no strains studied to date have observed interactions of nuclear transport pathways with nsp3. One possibility is the recruitment of nuclear transport proteins to 229E nsp3 to assist in the transport of viral components from double-membraned vesicles, which nsp3 is essential in helping to create^14–16^. A recent study showed that nsp3 from murine hepatitis C virus (MHV) not only assists in initial formation of the DMVs, but subsequently forms the core of a pore-like structure that may aid in export of viral RNA^34^. The absence of these protein interactions between nsp3.1 homologs from betacoronavirus strains and nuclear importins points to evolution of functional divergence.

We also find that MERS nsp3.2 uniquely interacts with rRNA processing machinery involved in ribosome biogenesis. Several of these factors (WDR36, NOL10, URB1) were previously found to interact with SARS-CoV-2, SARS-CoV, and MERS-CoV nsp8 homologs^18,19^. Some viruses, such as HIV-1, have also been shown to down-regulate factors involved in rRNA processing during infection^38^. Given the dependence of all coronaviruses on host translation machinery, it is interesting that MERS is the only CoV to maintain these interactions. This may represent a specialized avenue for MERS-CoV to modulate host translation through nsp3.

Viruses often modulate cellular stress responses in order to assist during replication and/or subvert the host immune system. Many groups of viruses, including coronaviruses and flaviviruses, have been shown to upregulate branches of the UPR^40,41,44,57–62^, although exact roles of this activation for the viral life cycle are still debated. We were therefore particularly interested to find ATF6 as a high confidence interactor of the SARS-CoV and SARS-CoV-2 nsp3.1 fragments. ATF6 is a transmembrane sensor representing one of the most upstream portions of the ER unfolded stress response (UPR). As noted, coronaviruses replicate near the ER, and nsp3 possesses transmembrane domains that anchor the protein in the ER membrane. Given the localization of nsp3.1 to the cytosol, the protein is likely to interact with the cytosolic basic leucine zipper (bZIP) transcriptional activator domain of ATF6. Therefore, we sought to determine if the fragment could be regulating the ability of ATF6 to respond to stress. We probed the transcriptional and translational level of known ATF6 regulated genes after overexpression of nsp3.1 fragments. Surprisingly, SARS-CoV-2 nsp3.1 led to a significant decrease in ATF6 protein markers, suggesting suppression of the ATF6 pathway. In addition to inhibiting basal ATF6 activity, we showed that SARS-CoV-2 nsp3.1 was also capable of blocking the activation of ATF6 by pharmacological activator **147**. It is likely that UPR activity has to be finely tuned during infection to prevent detrimental consequences from prolonged activation, such as apoptosis induction. Prior studies found that the IRE/XBP1s UPR branch was inhibited by the SARS-CoV E protein and that ATF6 gene targets were not upregulated during MHV infection despite ATF6 activation and cleavage^58,63^. These results suggest that coronaviruses employ strategies to attenuate distinct UPR signaling branches, but little is known about what viral proteins are responsible for tuning UPR activity. Our results indicate that the N-terminal regions of SARS-CoV-2 nsp3 directly acts on ATF6 to suppress activation. Further work is needed to understand the molecular mechanisms of this suppression and its role during SARS-CoV-2 infection.

Many studies have highlighted the utility of viral protein interactomes in identifying roles for host pathways in the life cycle of many viruses^36,64^, and large-scale interactome studies with SARS-CoV-2 demonstrated how these interactomes may be useful for identifying existing drugs to be repurposed to fight viral infections^18^. Despite the large body of work done in the past year on SARS-CoV-2, interactome studies of nsp3 have been limited to the PL2^pro^ papain-like-protease domain^11^. In addition to presenting data for the full nsp3 interactome, our studies highlight the utility of using tandem mass tag technology to quantitatively compare the SARS-CoV-2 nsp3 interactome to the interactome of other known coronaviruses, both severely and mildly pathogenic. We find very divergent interactomes, suggesting that while there is a large conservation of domains, there may also be more specific roles for this protein in the context of each individual virus. In the future, it may be important to study the variations in host protein interactions that occur between specific SARS-CoV-2 variants that are rapidly emerging. For instance, the recently isolated B.1.1.7 strain, shown to be highly transmissible and responsible for a large uptick in cases in the United Kingdom, possesses four mutations within orf1ab, three of which are in nsp3^65,66^. Additionally, the South Africa B.1.1.153 variant contains one nsp3 mutation, and the Brazilian P.1 variant contains two^65^. This makes nsp3 protein variants relevant to study for better understanding how such evolutionary adaptations in non-structural proteins may impact virulence.

## METHODS

### Construct design

The coding sequences for full-length nsp3 were obtained from GenBank (SARS-CoV-2 isolate Wuhan-Hu-1 MN908947, SARS-CoV Urbani AY279741, hCoV-OC43 NC006213, MERS-CoV JX869059, hCoV-229E AF304460).

Sequences for creating fragments of the SARS-CoV nsp3 were chosen based on Pan et al, 2008, PLoS One^21^. The amino acid sequences of full-length nsp3 for the remaining hCoV strains were aligned using ClustalOmega to the SARS-CoV fragments to determine the corresponding starting/ending positions for each fragment.

Full length SARS-CoV-2 nsp3 was codon optimized and cloned into a pcDNA-(+)-C-DYK vector (Genscript). Truncations were performed using primers listed in Table 1. All other nsp3 fragments were individually codon optimized and cloned into pTwist CMV Hygro vectors (Twist Biosciences). SARS-CoV and SARS-CoV-2 nsp3.1 strep-tagged constructs were created using primers listed in Table 1. Briefly, nsp3 fragment plasmids were amplified to exclude the FLAG tag. A pLVX vector containing a strep tag was used to amplify the insert. The two fragments were ligated using a DNA HiFi assembly kit (NEB). All plasmid constructs were confirmed by sequencing (Genewiz).

**Table 1.**
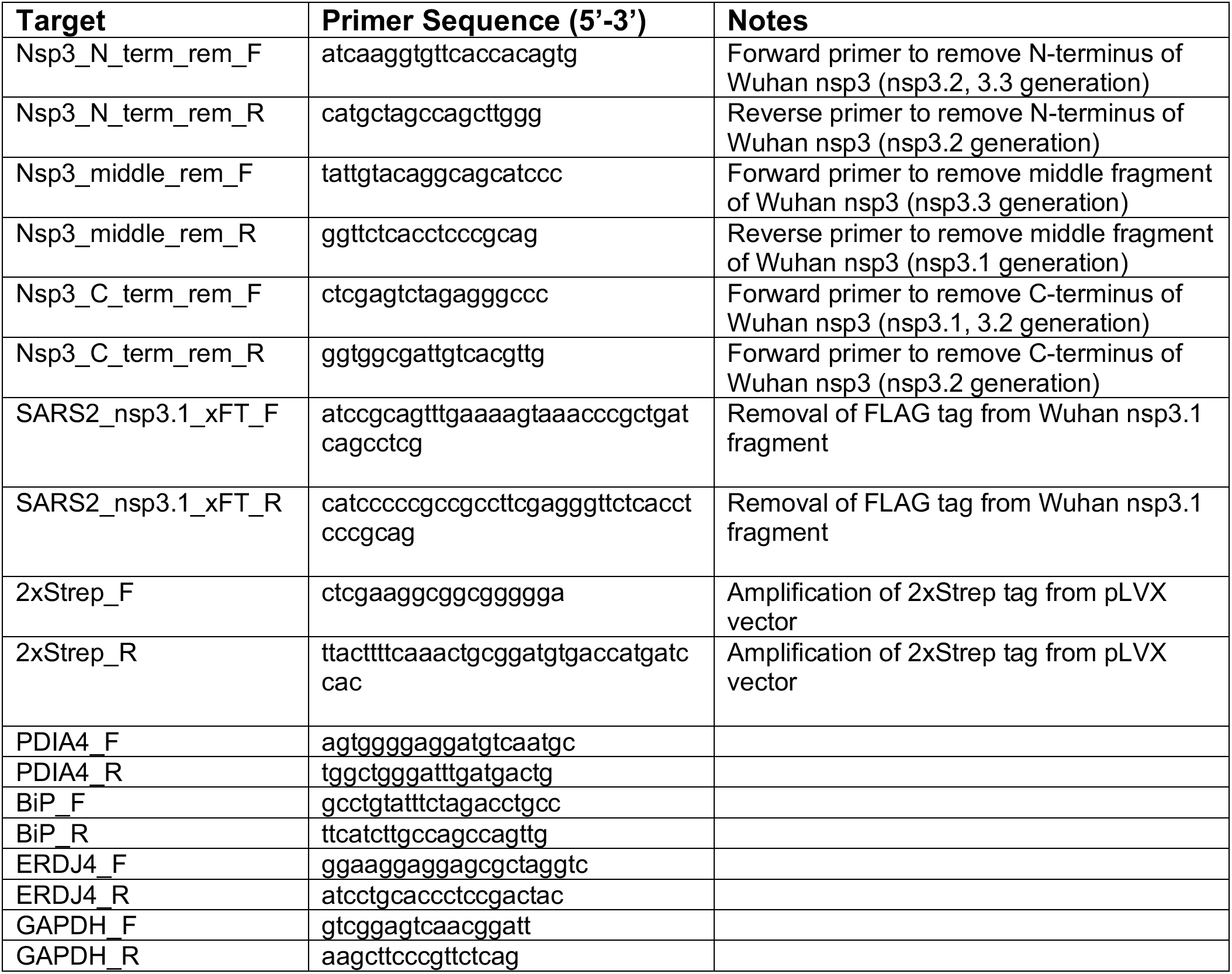
Primers used for qRT-PCR and generation of constructs

### Cell culture and transfection

HEK293T and HEK293T-REx cells were maintained in Dulbeccos’ Modified Eagle’s Medium (high glucose) and supplemented with 10% fetal bovine serum, 1% penicillin/streptomycin, and 1% glutamine. Cells were kept at 37°C, 5% CO_2_. For transfections, 2E6 cells were seeded into 10cm tissue culture dishes. 24 hours after seeding, cells were transfected using a calcium phosphate method with 5 μg nsp3 or tdTomato construct. Media was exchanged 16 hours post-transfection and cells were harvested 24 hours after media exchange.

### FLAG Immunoprecipitations

Immunoprecipitations were performed as reported previously^22^. Cells were collected from 10cm dishes via scraping, washed with PBS, and lysed by suspension in TNI buffer (50mM Tris pH 7.5, 150mM NaCl, 0.5% IGEPAL-CA-630) supplemented with Roche cOmplete protease inhibitor. Cells were left to lyse on ice for at least 10 minutes, followed by 10 minute sonication in a room temperature water bath. Lysates were cleared by centrifugation at 21.1xg for 20 minutes. Protein concentrations were normalized using 1x BioRad Protein Assay Dye, and normalized lysates were added to 15 μL pre-washed (4x in lysis buffer) Sepharose 4B beads (Sigma) and rocked at 4°C for 1 hour. Resin was collected by centrifugation at 400xg for 10 minutes, and pre-cleared supernatant was added to 15 μL G1 anti-DYKDDDDK resin (GenScript) and rocked at 4°C overnight. The next day, resin was collected by centrifugation at 400xg for 10 minutes. Resin was washed 4x with lysis buffer. Resin-bound proteins were eluted with the addition of modified 3x Laemelli buffer (6% SDS, 62.5mM Tris) for 30 minutes at room temperature, followed by 15 minutes at 37°C. A second elution was performed for 15 minutes at 37°C.

During the first IP for each construct, immunoprecipitation was confirmed by silver stain using a Pierce Silver Stain Kit (Thermo Fisher) and by western blotting with anti-FLAG antibody (Sigma-Aldrich, F1804)

### Nsp3.1-ST & ATF6-FT Co-immunoprecipitation

HEK293T-REx cells expressing doxycycline-inducible 6xHis-3xFLAG-HsATF6α were seeded in 15cm tissue culture dishes^67^. Cells were transiently transfected with 15 μg SARS-CoV-2 nsp3.1-ST or tdTomato constructs 24 hours after seeding as previously described. At 16 hours post-transfection media was exchanged and cells were treated with either DMSO or 100 μM doxycycline. Cells were harvested 24 hours later via scraping in cold PBS with 10 μM MG132. Cells were lysed in ATF6 lysis buffer (50 mM Tris pH 7.4, 150 mM NaCl, 5 mM EDTA, 1x protease inhibitor, and 1% LMNG (Anatrace NG322) for 15 min at 4°C as previously described^68^. Lysates were vortexed twice briefly and then cleared by centrifugation at 17,000 x g for 15 min. Lysates were normalized and pre-cleared with 15 μL Sepharose 4B beads in ATF6 wash buffer (50 mM Tris pH 7.4, 150 mM NaCl, 5 mM EDTA). Pre-cleared supernatant was then divided in two, with half immunoprecipitated for FLAG-ATF6 with 15 μL G1 anti-DYKDDDK resin and the other half affinity purified for nsp3.1-ST using 15 μL Strep-Tactin XT Superflow High Capacity resin (IBA Life Sciences, 2-4030-002) and rocked at 4°C overnight. Resin was washed 4x with ATF6 wash buffer. Resin-bound proteins were eluted with modified 3x Laemelli buffer (6% SDS, 62.5mM Tris) as described previously. Inputs and elutions were then diluted with 6x Laemelli buffer (12% SDS, 125 mM Tris pH 6.8, 20% glycerol, bromophenol blue, 100 mM DTT), heated at 37°C for 30 min, and run on SDS-PAGE gel. Proteins were detected via western blotting with M2 anti-FLAG (Sigma-Aldrich, F1804) and THE anti-Strep II tag FITC (Genescript, A01736-100) antibodies.

### Western blot and RT-qPCR analysis of UPR activation

HEK293T cells were transfected with nsp3.1-FT homologs or tdTomato (mock) in 6-well plates as previously described. Mock samples were treated with DMSO or 6 mg/mL Tunicamycin (Tm) for 16 h (protein analysis) or 6 h (transcript analysis) prior to harvest by cell scraping. Harvested cells were split into a 2/3 aliquot for protein analysis and a 1/3 aliquot for mRNA analysis.

For protein analysis, cells were lysed as previously described for immunoprecipitations. Lysates were normalized using 1x BioRad Protein Assay Dye and run on SDS-PAGE gel before transfer to PVDF membranes and subsequent western blotting. Blots were probed with 1:1000 dilutions of anti-KDEL (Enzo, ADI-SPA-827-F), anti-PDIA4 (ProteinTech, 14712-1-AP), M2 anti-FLAG (Sigma-Aldrich, F1804), and anti-GAPDH (GeneTex, GTX627408) antibodies in 5% BSA.

For RT-qPCR, cellular RNA was extracted using the Zymo Quick-RNA miniprep kit. Then 500 ng total cellular RNA was synthesized into cDNA using random hexamer primers (IDT), oligo-dT primers (IDT), and Promega M-MLV reverse transcriptase. Subsequent qPCR analysis was carried out using BioRad iTaq Universal SYBR Green Supermix, combined with the primers listed below for target genes. Reactions were run in 96-well plates on a BioRad CFX qPCR instrument. Conditions used for amplification were 95°C, 2 minutes, 45 repeats of 95°C, 10s and 60°C, 30s. A melting curve was generated in 0.5 °C intervals from 65 °C to 95 °C. Cq values were calculated by the BioRad CFX Maestro software. Transcripts were normalized to a housekeeping gene (GAPDH). All measurements were performed in technical duplicate; each of these duplicates was treated as a single measurement for the final average. Data was analyzed using the BioRad CFX Maestro software.

### Global proteomics of UPR activation

HEK293T cells were transfected with SARS-CoV-2 nsp3.1-FT or tdTomato (mock) in 10 cm dishes in triplicate as previously described. Samples were treated with either DMSO or 10 μM **147** for 16 h prior to harvest by cell scraping. Harvested cells were lysed in TNI lysis buffer and 10 μg of protein (as measured using Bio-Rad protein assay dye reagent) was aliquoted and precipitated via methanol/chloroform precipitation as described below. Pellets were digested with trypsin (Thermo Fisher) and labeled with 16plex TMTpro.

### Tandem Mass Tag sample preparation

Sample preparation was carried out as previously described^22^. Eluted proteins were precipitated via methanol/chloroform water (3:1:3) and washed thrice with methanol. Each wash was followed by a 5 minute spin at 10,000xg. Protein pellets were air dried and resuspended in 1% Rapigest SF (Waters). Resuspended proteins were reduced (TCEP) for 30 minutes, alkylated (iodoacetamide) for 30 minutes, and digested with 0.5 μg trypsin/Lys-C (Thermo Fisher) overnight. Digested peptides were labeled using 16plex TMTpro (Thermo Scientific) and quenched with the addition of ammonium bicarbonate. Samples were pooled, acidified, and concentrated. Cleaved Rapigest products were removed by centrifugation at 17,000xg for 45 minutes.

### MudPIT LC-MS/MS analysis

Triphasic MudPIT columns were prepared as previously described using alternating layers of 1.5cm C18 resin, 1.5cm SCX resin, and 1.5cm C18 resin^69^. Pooled TMT samples (roughly one-third of pooled IP samples and 10 μg of peptide from global UPR activation samples) were loaded onto the microcapillaries using a high-pressure chamber, followed by a 30 minute wash in buffer A (95% water, 5% acetonitrile, 0.1% formic acid). Peptides were fractionated online by liquid chromatography using an Ultimate 3000 nanoLC system and subsequently analyzed using an Exploris480 mass spectrometer (Thermo Fisher). The MudPIT columns were installed on the LC column switching valve and followed by a 20cm fused silica microcapillary column filled with Aqua C18, 3μm, C18 resin (Phenomenex) ending in a laser-pulled tip. Prior to use, columns were washed in the same way as the MudPIT capillaries. MudPIT runs were carried out by 10μL sequential injections of 0, 10, 20, 40, 60, 80, 100 % buffer C (500mM ammonium acetate, 94.9% water, 5% acetonitrile, 0.1% formic acid) for IP samples and 0, 10, 20, 30, 40, 50, 60, 70, 80, 90, 100% buffer C for global UPR activation samples, followed by a final injection of 90% C, 10% buffer B (99.9% acetonitrile, 0.1% formic acid v/v). Each injection was followed by a 130 min gradient using a flow rate of 500nL/min (0-6 min: 2% buffer B, 8 min: 5% B, 100 min: 35% B, 105min: 65% B, 106-113 min: 85% B, 113-130 min: 2% B). ESI was performed directly from the tip of the microcapillary column using a spray voltage of 2.2 kV, an ion transfer tube temperature of 275°C and a RF Lens of 40%. MS1 spectra were collected using a scan range of 400-1600 m/z, 120k resolution, AGC target of 300%, and automatic injection times. Data-dependent MS2 spectra were obtained using a monoisotopic peak selection mode: peptide, including charge state 2-7, TopSpeed method (3s cycle time), isolation window 0.4 m/z, HCD fragmentation using a normalized collision energy of 32 (TMTpro), resolution 45k, AGC target of 200%, automatic injection times, and a dynamic exclusion (20 ppm window) set to 60s.

### Data analysis

nsp3.1 analysis combined 5 individual MS runs totaling 62 samples (15x tdTomato, 13x SARS-CoV-2, 9x SARS-CoV, 8x MERS-CoV, 9x hCoV-OC43, 8x hCoV-229E). nsp3.3 analysis combined 5 individual MS runs totaling 63 samples (15x tdTomato, 14x SARS-CoV-2, 9x SARS-CoV, 8x MERS-CoV, 9x hCoV-OC43, 8x hCoV-229E). nsp3.2 analysis combined 4 individual MS runs totaling 63 co-IP samples (tdTomato 15, SARS-CoV-2 14, SARS-CoV 9, hCoV-OC43 9, MERS-CoV 8, hCoV-229E 8). Identification and quantification of peptides were performed in Proteome Discoverer 2.4 (Thermo Fisher) using the SwissProt human database (TaxID 9606, released 11/23/2019) with nsp3 fragment sequences manually added. Searches were conducted with Sequest HT using the following parameters: trypsin cleavage (maximum 2 missed cleavages), minimum peptide length 6 AAs, precursor mass tolerance 20ppm, fragment mass tolerance 0.02Da, dynamic modifications of Met oxidation (+15.995Da), protein N-terminal Met loss (−131.040Da), and protein N-terminal acetylation (+42.011Da), static modifications of TMTpro (+304.207Da) at Lys and N-termini and Cys carbamidomethylation (+57.021Da). Peptide IDs were filtered using Percolator with an FDR target of 0.01. Proteins were filtered based on a 0.01 FDR, and protein groups were created according to a strict parsimony principle. TMT reporter ions were quantified considering unique and razor peptides, excluding peptides with co-isolation interference greater that 25%. Peptide abundances were normalized based on total peptide amounts in each channel, assuming similar levels of background in the IPs. For global UPR proteomics, protein abundances were also scaled. Protein quantification used all quantified peptides. Post-search filtering was done to include only proteins with two identified peptides. Pairwise ratios between conditions were calculated in Proteome Discoverer based on total protein abundance, and ANOVA was performed on individual proteins to test for change in abundances and report adjusted P-values. To filter interactors of individual nsp3 fragments, we used a variable cutoff method combining log2 enrichment and adjusted p-value according to a published method^22,70^. The histogram of log2 protein abundance fold changes for each construct vs the tdTomato control was fitted to a gaussian curve with a bin width of 0.1 using a nonlinear least square fit (excluding outliers) to determine the standard deviation σ of the scatter. For medium- and high-confidence interactors, the cutoff values were 1σ and 2σ respectively. To take into consideration the adjusted p-values, we used a hyperbolic curve *y > c(x-x_0_)* where y is the adjusted p-value, x is the log2 fold change, and x_0_ corresponds to the value of the 1σ or 2σ standard deviation.

### Geneset enrichment analysis, comparative heatmaps, and network plots

A gene ontology (GO) enrichment analysis for biological processes was conducted in EnrichR. The analysis was conducted separately for the high (and/or medium) confidence interactors for each fragment. GO terms were manually filtered for adjusted p-values <0.1. Redundant GO terms were grouped manually based on overlapping genes in related pathways.

Network plots were generated in Cytoscape^71^; human protein interactions were validated based on the STRING database.

## ACKNOWLEDGEMENTS

We thank Andrew Spencer, Joyce Kariuki, and Athira Sivadas for assistance with data analysis and literature review, as well as members of the Plate lab for their critical reading and feedback of this manuscript. This work was funded by T32 AI112451 (NIAID) and National Science Foundation Graduate Research Fellowship (KMA); T32 GM008554 (NIGMS) (JPD); 1R35GM133552 (NIGMS); and Vanderbilt University funds.

## DATA AVAILABILITY

The mass spectrometry proteomics data have been deposited to the ProteomeXchange Consortium via the PRIDE partner repository with the dataset identifier PXD024566. All other necessary data are contained within the manuscript.

