## Supporting Information for "Comparative host interactomes of the SARS-CoV-2 nonstructural protein 3 and human coronavirus homologs"

### TABLE OF CONTENT

Figures S1 – S13 pp S2-S16

Tables S1 – S8 p S17

Supporting Information References pp S18-S19

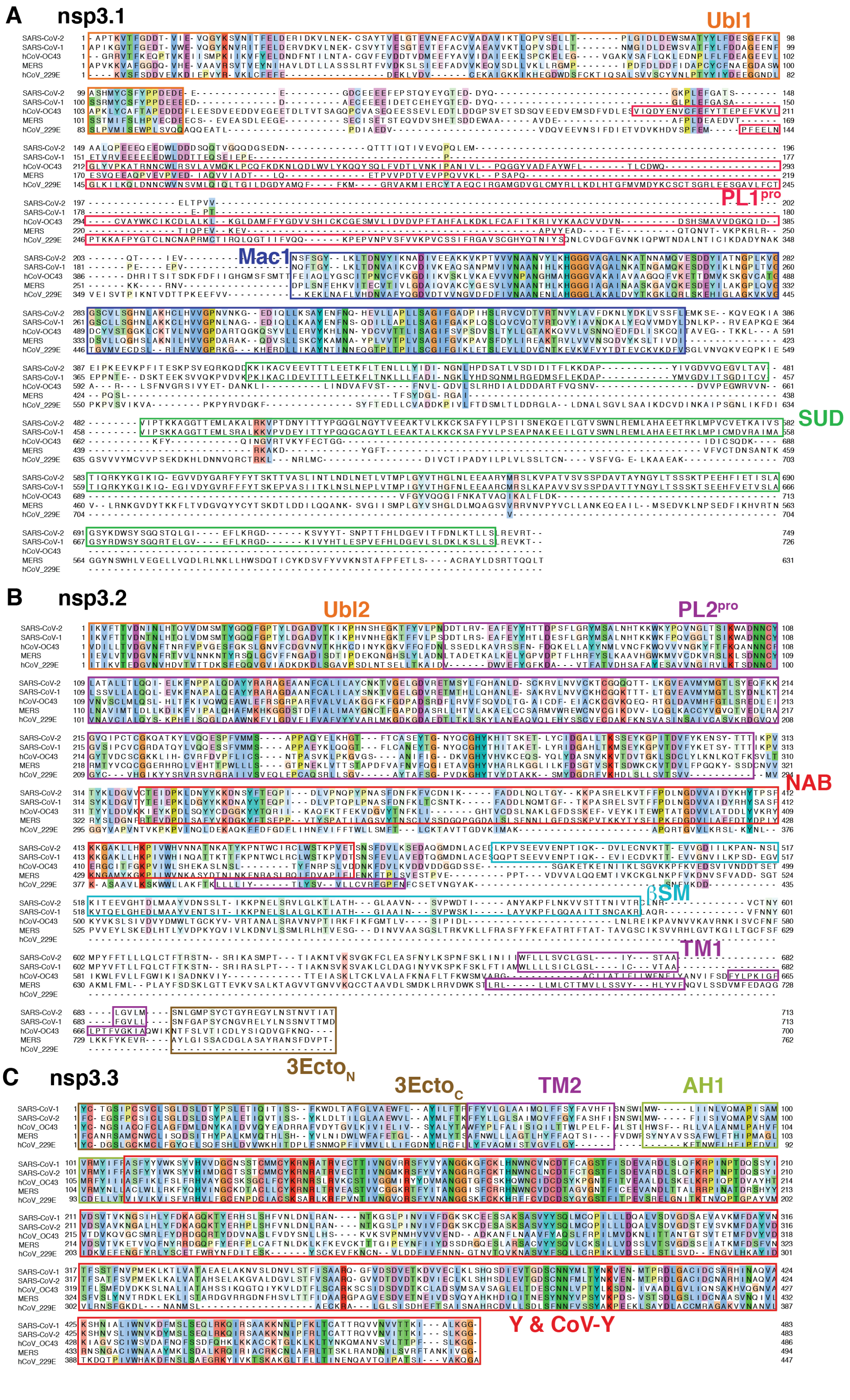
A nsp3.1
Ubl1
PL1pro
Mac1
SUD
B nsp3.2
Ubl2
PL2pro
NAB
βSM
TM1
3EctoN
C nsp3.3
3EctoC
TM2
AH1
Y & CoV-Y
SARS-CoV-2
SARS-CoV-1
hCoV-OC43
MERS
hCoV_229E

**Figure S1. Multiple-sequence alignment and domain organization of amino acid sequences for nsp3 truncations from different coronavirus strains.**

**A-C.** Amino acid sequences of nsp3.1 (A), nsp3.2 (B), and nsp3.3 (C) homologs were aligned using Clustal Omega. Domain organization is noted by colored boxes[1].

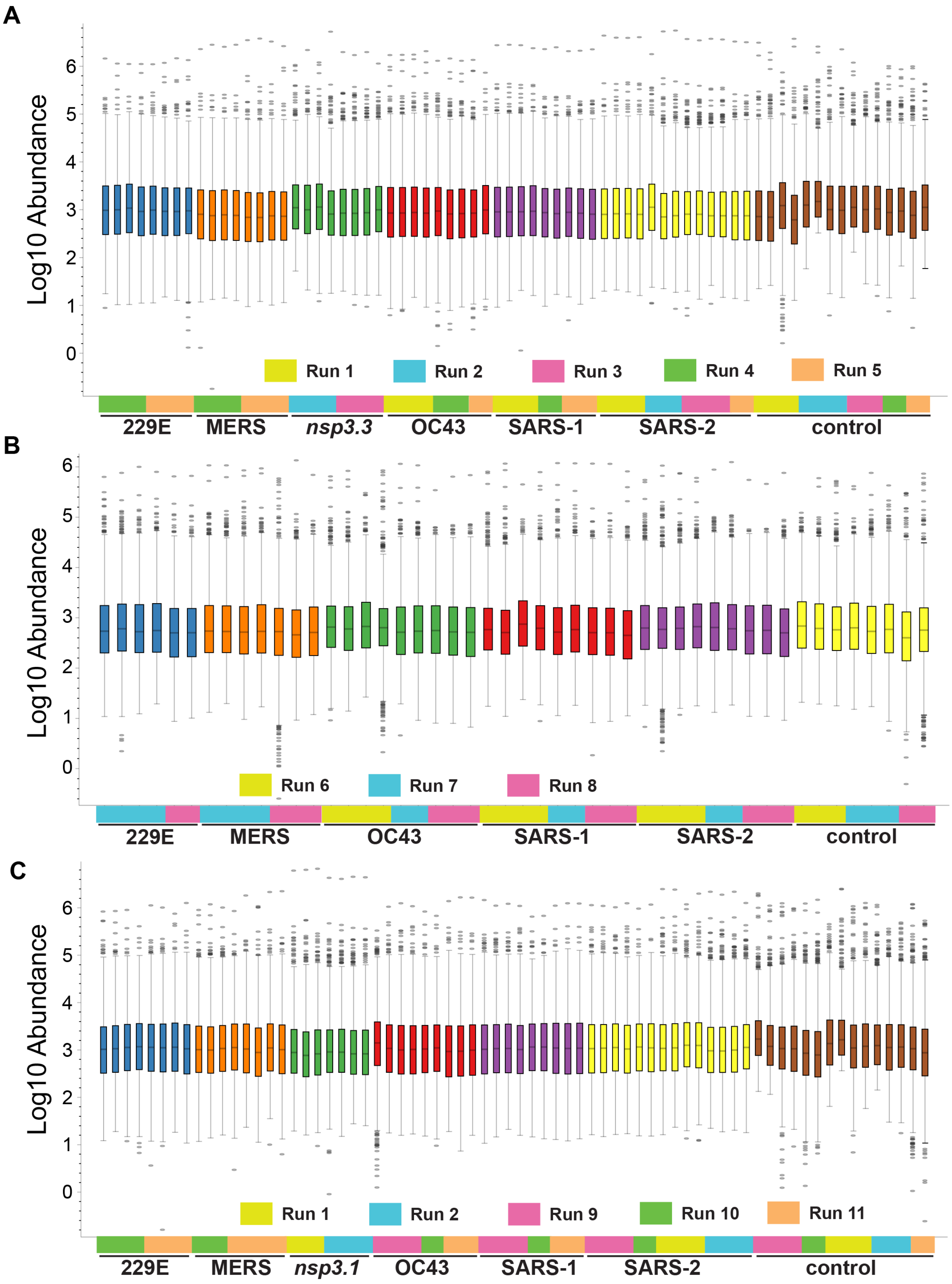
A
Log10 Abundance
0
1
2
3
4
5
6
Run 1
Run 2
Run 3
Run 4
Run 5
229E
MERS
nsp3.3
OC43
SARS-1
SARS-2
control
B
Log10 Abundance
Run 6
Run 7
Run 8
229E
MERS
OC43
SARS-1
SARS-2
control
C
Log10 Abundance
Run 1
Run 2
Run 9
Run 10
Run 11
229E
MERS
nsp3.1
OC43
SARS-1
SARS-2
control

**Figure S2. Nsp3 TMT intensity distribution.**

**A-C.** Box-and-whisker plot of the log10 TMT intensity abundance of all 11 mass spectrometry runs used in this study. Nsp3 homologs are grouped by same color box-and-whiskers while the identity of the corresponding mass spectrometry run is denoted by colored blocks below. TMT abundances were normalized based on total peptide amounts. See supporting dataset S8 for the layout of samples across TMT channels.

**A.** Normalized TMT abundance distribution for nsp3.1 dataset. Channels denoted "*nsp3.3*" were used for the nsp3.3 dataset within the corresponding mass spectrometry run.

**B.** Normalized TMT abundance distribution for nsp3.2 dataset.

**C.** Normalized TMT abundance distribution for nsp3.3 dataset. Channels denoted "*nsp3.1*" were used for the nsp3.1 dataset within the corresponding mass spectrometry run.

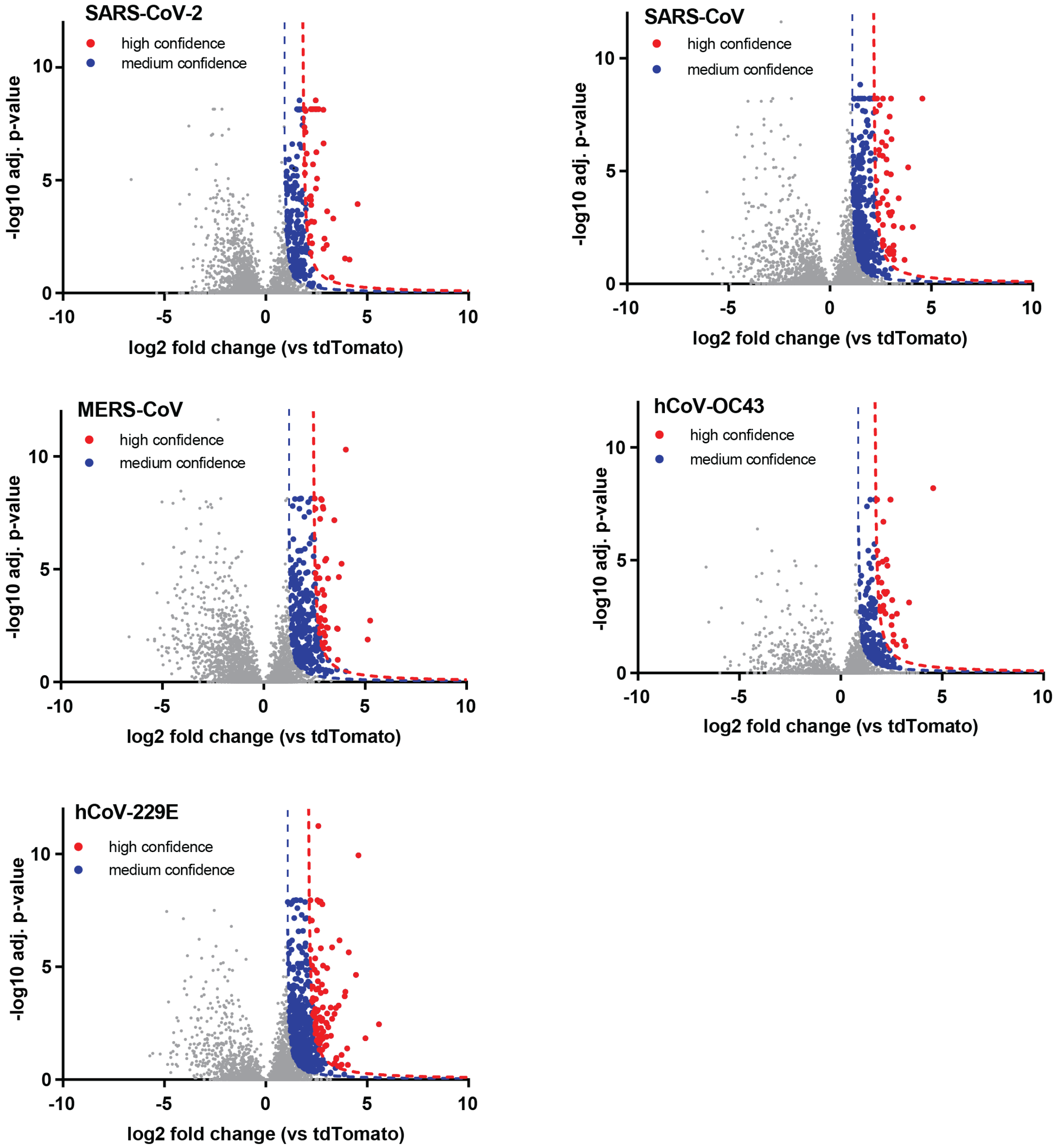


**Figure S3. Volcano plots of nsp3.1 homolog high-confidence interactors enriched vs tdTomato control.**

Host interactors of nsp3.1 homologs were identified by quantitative proteomics and graphed by log2 fold change compared to tdTomato control and -log10 adjusted *p*-value (based on ANOVA). Interactors were filtered for medium (blue) and high (red) confidence interactors based on a hyperbolic curve using 1σ (medium-confidence) or 2σ (high-confidence) standard deviations of the histogram of log2 protein abundance fold changes (refer to “Data analysis” in Methods).

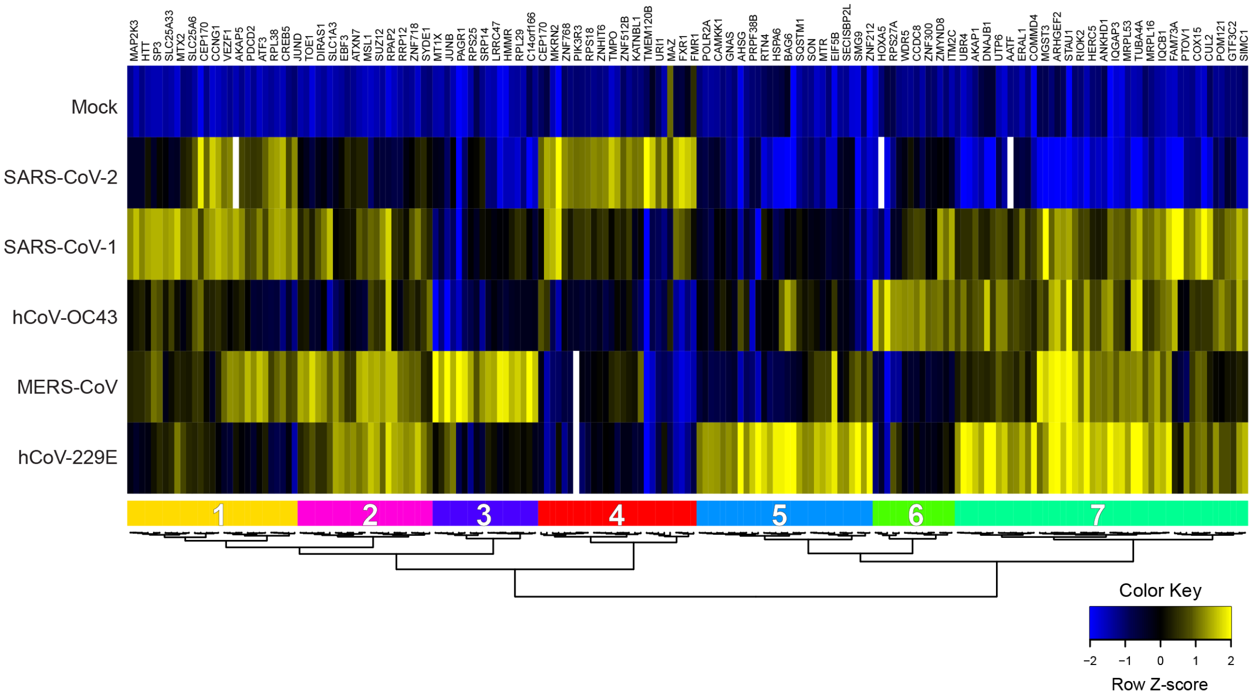


**Figure S4. Comparative heatmap of nsp3.1 high-confidence interactors.**
Unbiased hierarchical clustering of log2 fold change for high-confidence interactors of nsp3.1 homologs yields 7 unique clusters.

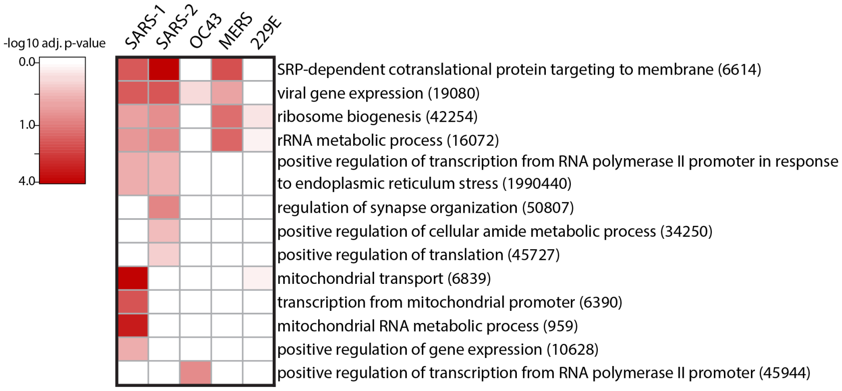


**Figure S5. GO term analysis of nsp3.1 high-confidence interactors.**
Heatmap of gene ontology (GO) term analysis of nsp3.1 high-confidence interactors for enriched biological processes. Selected terms are displayed with corresponding -log10 adjusted *p*-value for each homolog.

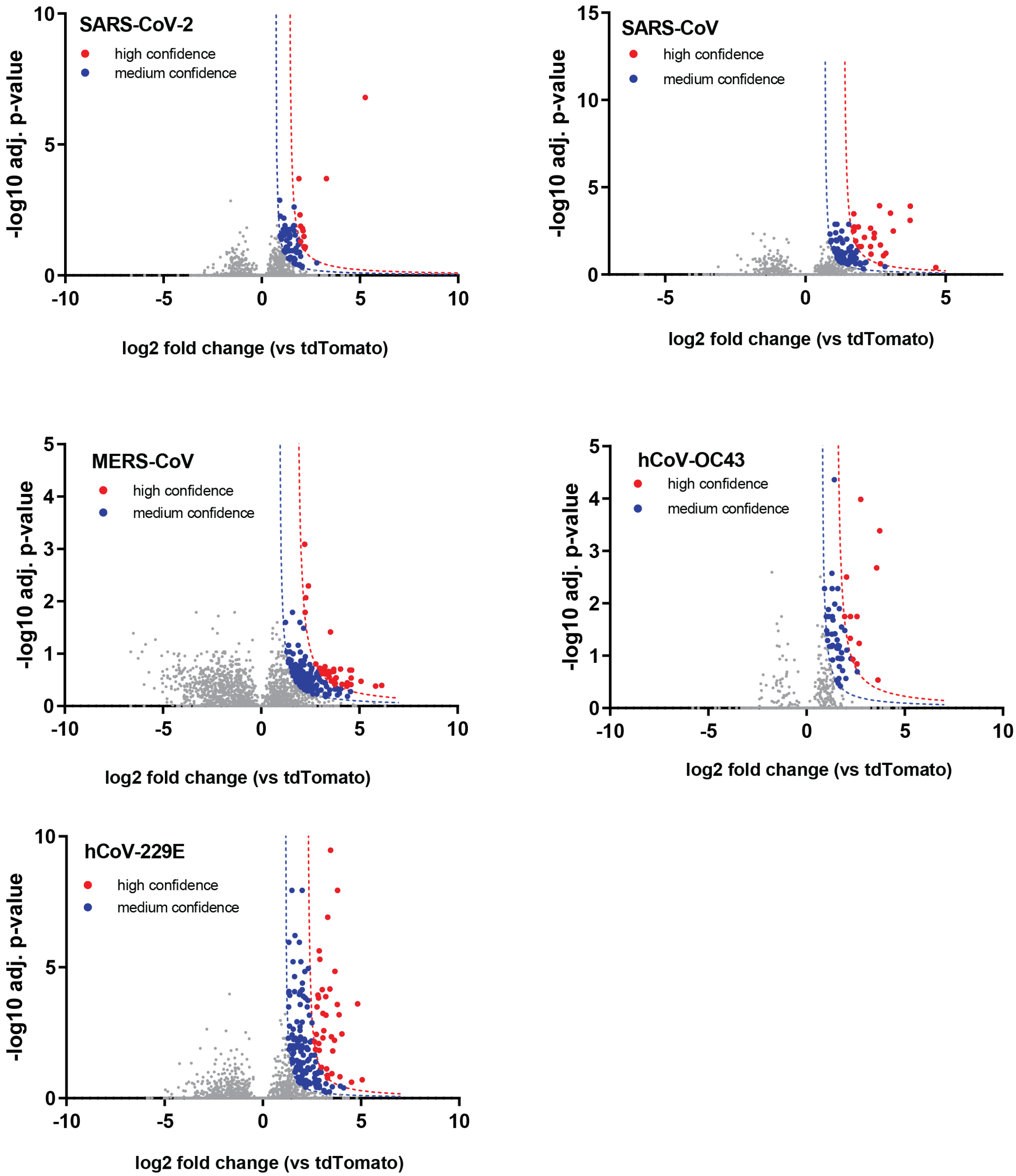


**Figure S6. Volcano plots of nsp3.2 homolog high-confidence interactors enriched vs tdTomato control.**

Host interactors of nsp3.2 homologs were identified by quantitative proteomics and graphed by log2 fold change compared to tdTomato control and -log10 adjusted *p*-value (based on ANOVA). Interactors were filtered for medium (blue) and high (red) confidence interactors based on a hyperbolic curve using 1σ (medium-confidence) or 2σ (high-confidence) standard deviations of the histogram of log2 protein abundance fold changes (refer to “Data analysis” in Methods).

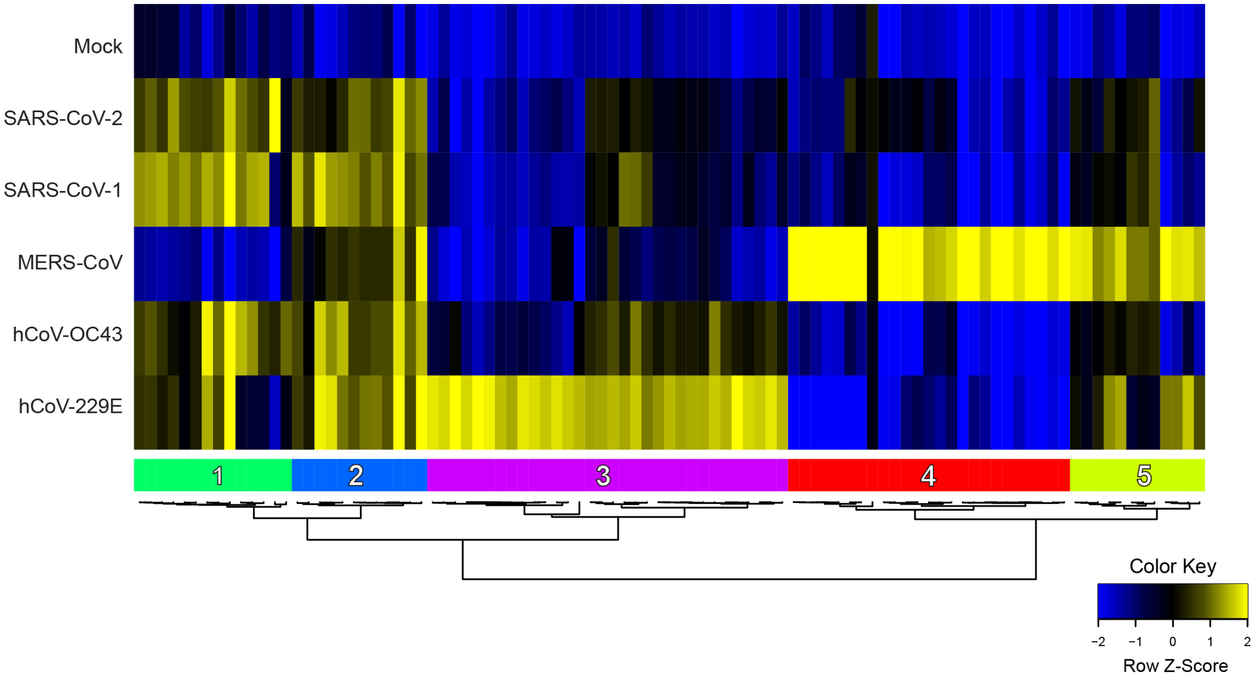


**Figure S7. Comparative heatmap of nsp3.2 high-confidence interactors.**
Unbiased hierarchical clustering of log2 fold change for high-confidence interactors of nsp3.2 homologs yields 5 unique clusters.

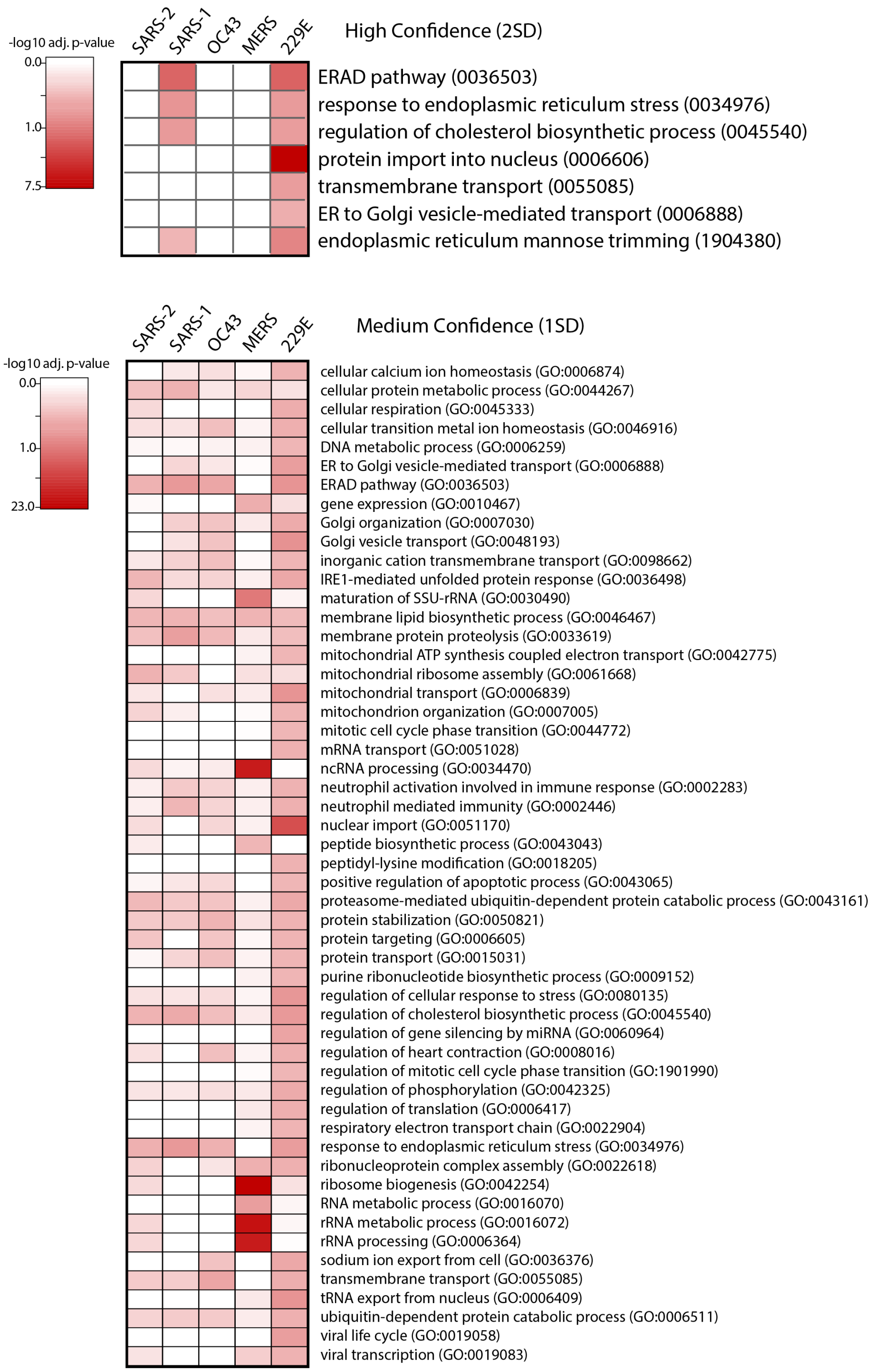


**Figure S8. GO term analysis of nsp3.2 high-confidence interactors.**
Heatmap of gene ontology (GO) term analysis of nsp3.2 high-confidence or medium-confidence interactors for enriched biological processes. Selected terms are displayed with corresponding -log10 adjusted *p*-value for each homolog.

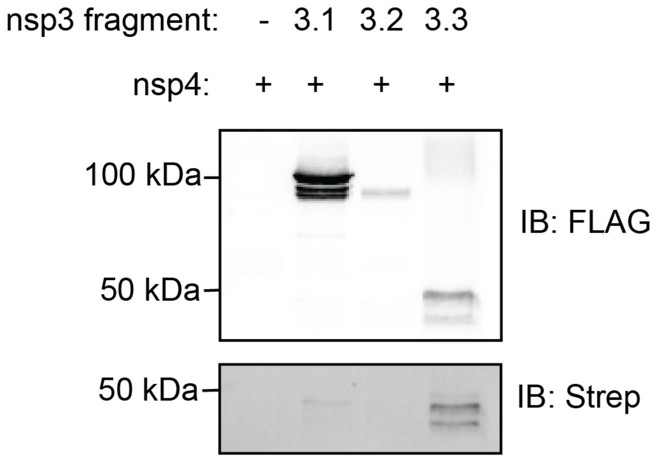


**Figure S9. Nsp3 fragment co-immunoprecipitations with nsp4.**
Samples were co-transfected with 2x-strep tagged SARS-CoV-2 nsp4[2] and individual SARS-CoV-2 FLAG-tagged nsp3 fragments (nsp3.1, nsp3.2, nsp3.3) or tdTomato as a control, respectively. FLAG immunoprecipitations were performed to pull down on nsp3 constructs and immunoblotting was performed with an anti-FLAG antibody to confirm nsp3.3 expression and an FITC anti-Strep antibody to monitor nsp4 co-immunoprecipitation.

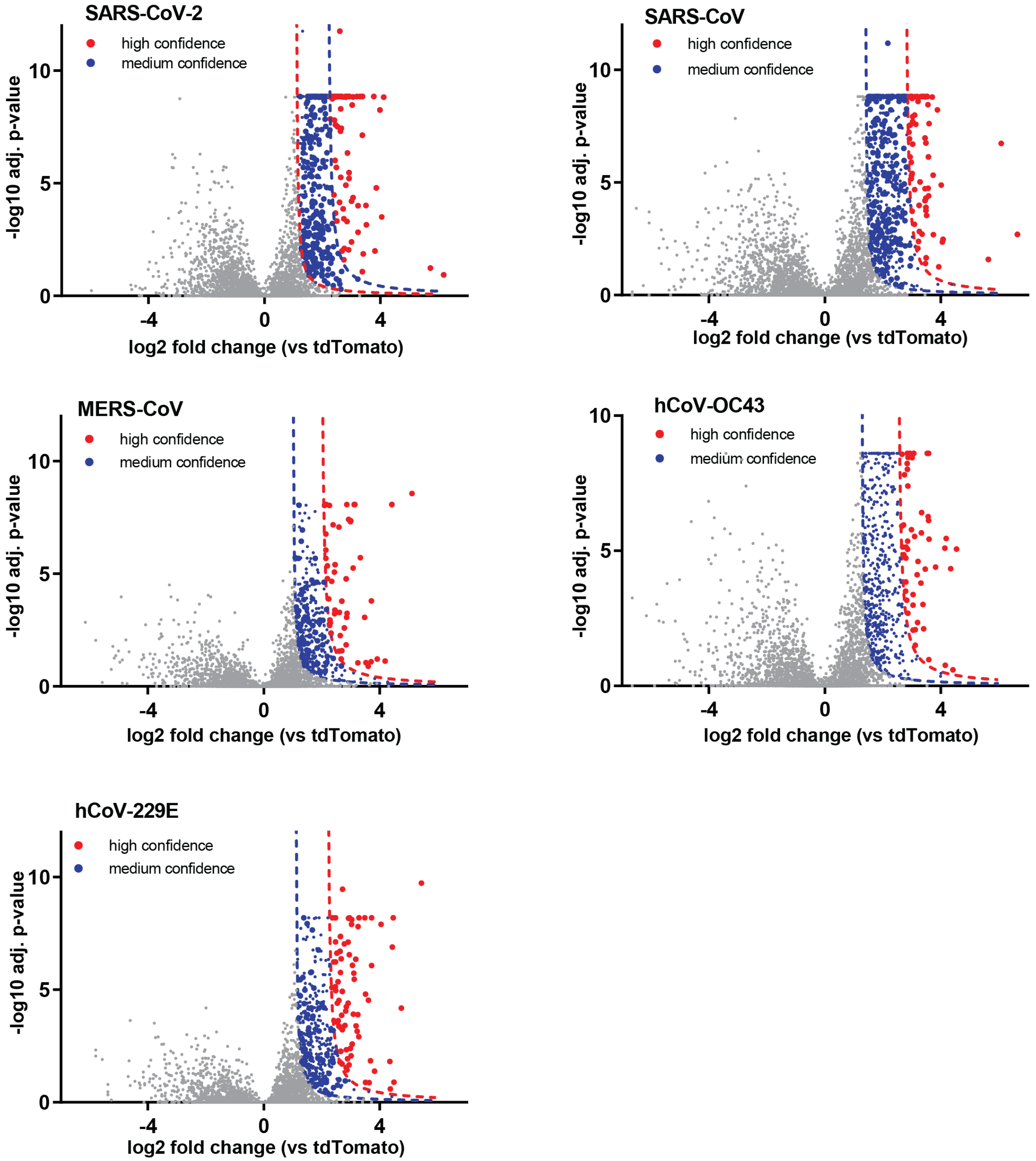


**Figure S10. Volcano plots of nsp3.3 homolog high-confidence interactors enriched vs tdTomato control.**

Host interactors of nsp3.3 homologs were identified by quantitative proteomics and graphed by log2 fold change compared to tdTomato control and -log10 adjusted *p*-value (based on ANOVA). Interactors were filtered for medium (blue) and high (red) confidence interactors based on a hyperbolic curve using 1σ (medium-confidence) or 2σ (high-confidence) standard deviations of the histogram of log2 protein abundance fold changes (refer to “Data analysis” in Methods).

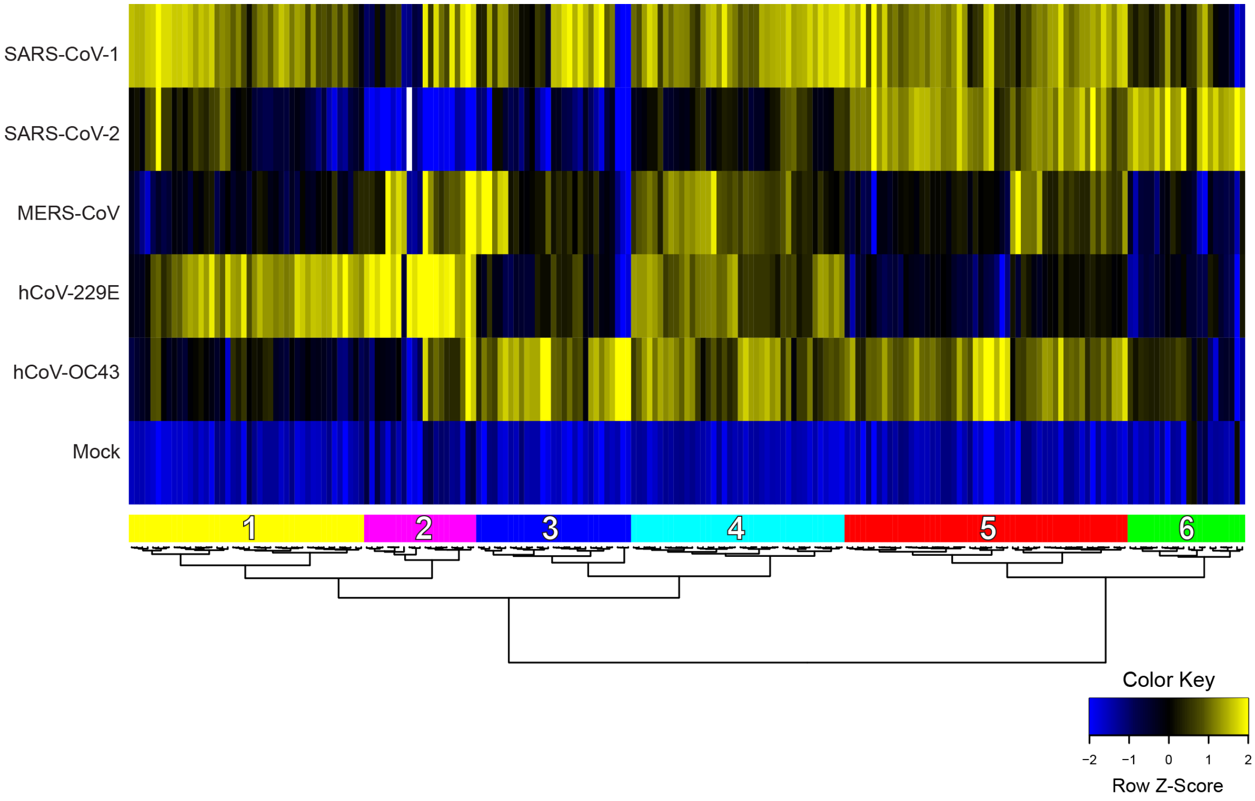


**Figure S11. Comparative heatmap of nsp3.3 high-confidence interactors.**
Unbiased hierarchical clustering of log2 fold change for high-confidence interactors of nsp3.3 homologs yields 6 unique clusters.

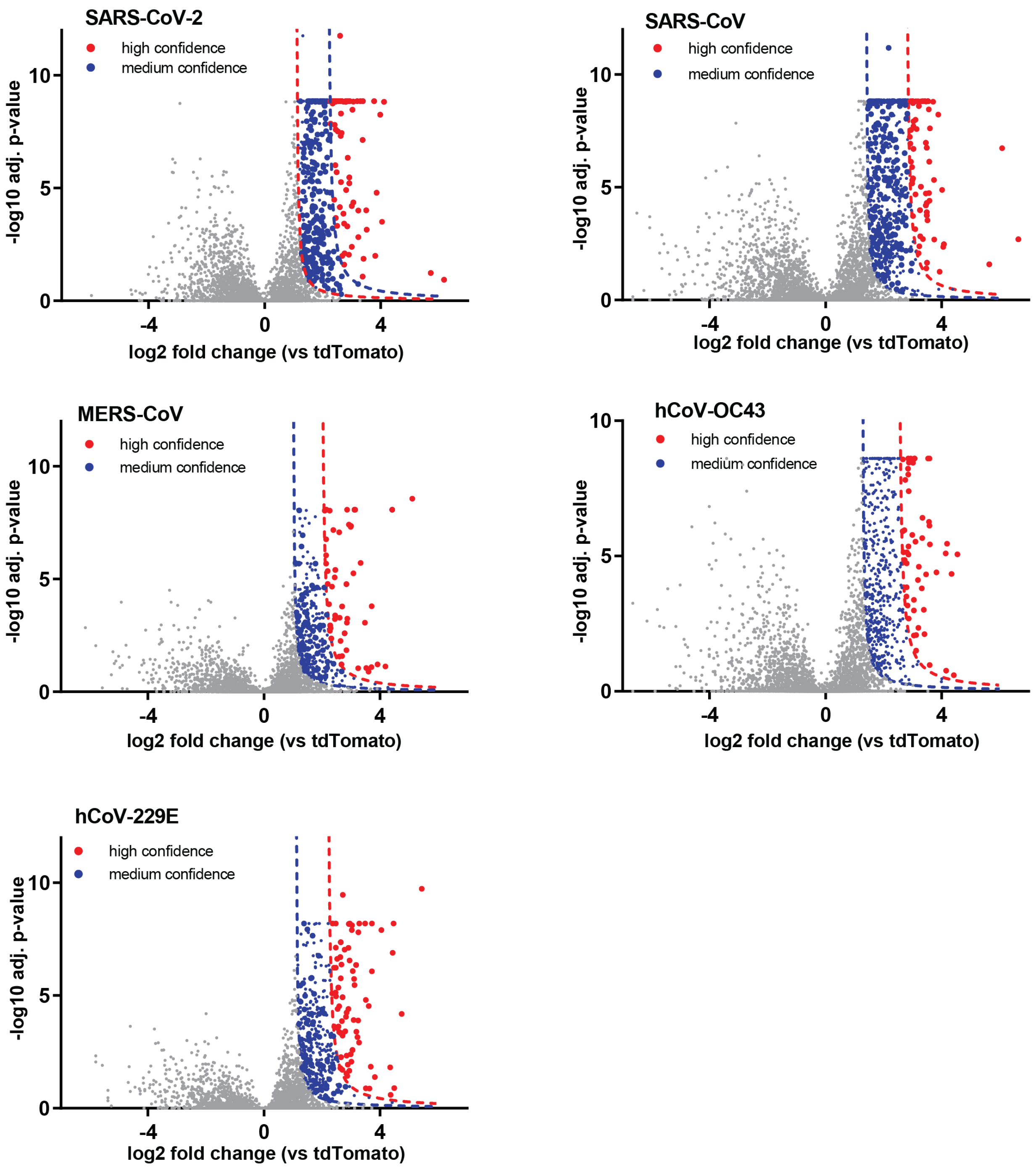


**Figure S12. GO term analysis of nsp3.3 high-confidence interactors.**
Heatmap of gene ontology (GO) term analysis of nsp3.3 high-confidence or medium-confidence interactors for enriched biological processes. Selected terms are displayed with corresponding -log10 adjusted *p*-value for each homolog.

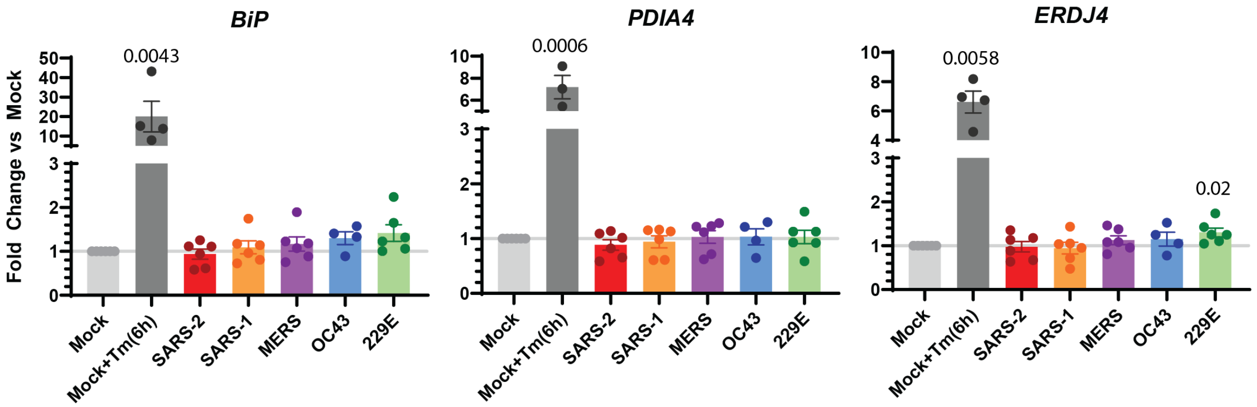


**Figure S13. Effect of nsp3.1 homologs on UPR transcript expression.**
Expression changes of UPR marker transcripts in response to nsp3.1 homolog or mock (tdTomato) transfection measured by reverse transcription qPCR (RT-qPCR). A mock transfected sample was treated with 6 µg/mL Tunicamycin (Tm) 6 h pre-harvest to induce general UPR activation as a positive control. *BiP* and *PDIA4* were measured as ATF6 markers, while *ERDJ4* was measured as a IRE1/XBP1s marker. Bars indicate average, error bars indicate SEM. Paired student T-tests were used to test for significance, with *p*-values < 0.05 shown. Mock+Tm(6h), *n* = 4 biological replicates; all other samples, *n* = 6 biological replicates.

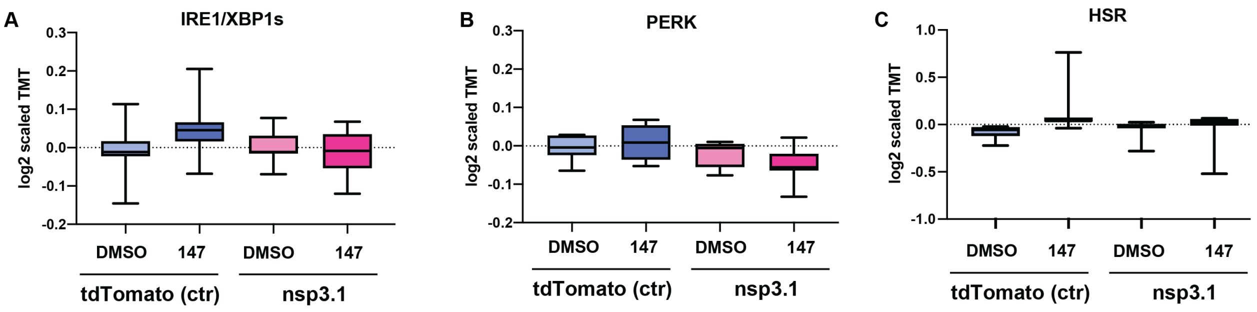


**Figure S14. IRE1 and PERK UPR activation by SARS-CoV-2 nsp3.1**

**A-B.** Box-and-whisker plots showing distribution of protein makers of IRE1/XBP1s (**A**) and PERK (**B**) UPR signaling branches as measured by quantitative proteomics. HEK293T cells were transfected with tdTomato (control) or SARS-CoV-2 nsp3.1-FT. Samples were treated with DMSO or 10 µM **147** for 16 h pre-harvest. A one-way ANOVA with Geisser-Greenhouse correction and post-hoc Tukey's multiple comparison test was used to determine significance (adjusted *p*-values all > 0.05). *n* = 3 biological replicates in a single mass spectrometry run.

### SUPPORTING DATASETS

(all supporting tables are available as separate Excel files)

**Table S1.** Proteomics data of comparative nsp3.1 interactome profiling. Included are protein identifications, quantifications, abundance ratios, statistical analysis, and filtering of medium- and high-confidence interactors.

**Table S2.** List of peptide identifications, quantifications, and protein mapping for comparative nsp3.1 interactome profiling.

**Table S3.** Proteomics data of comparative nsp3.2 interactome profiling. Included are protein identifications, quantifications, abundance ratios, statistical analysis, and filtering of medium- and high-confidence interactors.

**Table S4.** List of peptide identifications, quantifications, and protein mapping for comparative nsp3.2 interactome profiling.

**Table S5.** Proteomics data of comparative nsp3.3 interactome profiling. Included are protein identifications, quantifications, abundance ratios, statistical analysis, and filtering of medium- and high-confidence interactors.

**Table S6.** List of peptide identifications, quantifications, and protein mapping for comparative nsp3.3 interactome profiling.

**Table S7.** Global proteomics data of nsp3.1 modulation of the UPR and associated pathways. Included are protein identifications, quantifications, abundance ratios, statistical analysis, and filtering of UPR and related pathway markers.

**Table S8.** TMT channel organization for all quantitative proteomics experiments.

### SUPPORTING INFORMATION REFERENCES


(1) Lei, J.; Kusov, Y.; Hilgenfeld, R. Nsp3 of Coronaviruses: Structures and Functions of a Large Multi-Domain Protein. *Antiviral Research*. Elsevier B.V. January 1, 2018, pp 58–74. https://doi.org/10.1016/j.antiviral.2017.11.001.

(2) Gordon, D. E.; Jang, G. M.; Bouhaddou, M.; Xu, J.; Obernier, K.; White, K. M.; O'Meara, M. J.; Rezelj, V. V.; Guo, J. Z.; Swaney, D. L.; Tummino, T. A.; Hüttenhain, R.; Kaake, R. M.; Richards, A. L.; Tutuncuoglu, B.; Foussard, H.; Batra, J.; Haas, K.; Modak, M.; Kim, M.; Haas, P.; Polacco, B. J.; Braberg, H.; Fabius, J. M.; Eckhardt, M.; Soucheray, M.; Bennett, M. J.; Cakir, M.; McGregor, M. J.; Li, Q.; Meyer, B.; Roesch, F.; Vallet, T.; Mac Kain, A.; Miorin, L.; Moreno, E.; Naing, Z. Z. C.; Zhou, Y.; Peng, S.; Shi, Y.; Zhang, Z.; Shen, W.; Kirby, I. T.; Melnyk, J. E.; Chorba, J. S.; Lou, K.; Dai, S. A.; Barrio-Hernandez, I.; Memon, D.; Hernandez-Armenta, C.; Lyu, J.; Mathy, C. J. P.; Perica, T.; Pilla, K. B.; Ganesan, S. J.; Saltzberg, D. J.; Rakesh, R.; Liu, X.; Rosenthal, S. B.; Calviello, L.; Venkataramanan, S.; Liboy-Lugo, J.; Lin, Y.; Huang, X. P.; Liu, Y. F.; Wankowicz, S. A.; Bohn, M.; Safari, M.; Ugur, F. S.; Koh, C.; Savar, N. S.; Tran, Q. D.; Shengjuler, D.; Fletcher, S. J.; O'Neal, M. C.; Cai, Y.; Chang, J. C. J.; Broadhurst, D. J.; Klippsten, S.; Sharp, P. P.; Wenzell, N. A.; Kuzuoglu-Ozturk, D.; Wang, H. Y.; Trenker, R.; Young, J. M.; Cavero, D. A.; Hiatt, J.; Roth, T. L.; Rathore, U.; Subramanian, A.; Noack, J.; Hubert, M.; Stroud, R. M.; Frankel, A. D.; Rosenberg, O. S.; Verba, K. A.; Agard, D. A.; Ott, M.; Emerman, M.; Jura, N.; von Zastrow, M.; Verdin, E.; Ashworth, A.; Schwartz, O.; d'Enfert, C.; Mukherjee, S.; Jacobson, M.; Malik, H. S.; Fujimori, D. G.; Ideker, T.; Craik, C. S.; Floor, S. N.; Fraser, J. S.; Gross, J. D.; Sali, A.; Roth, B. L.; Ruggero, D.; Taunton, J.; Kortemme, T.; Beltrao, P.; Vignuzzi, M.; García-Sastre, A.; Shokat, K. M.; Shoichet, B. K.; Krogan, N. J. A SARS-CoV-2 Protein Interaction Map Reveals Targets for Drug Repurposing. *Nature* **2020**, *583* (7816), 459–468. https://doi.org/10.1038/s41586-020-

2286-9.
